# Genetic reprogramming by brief inhibition of the renin-angiotensin system in spontaneously hypertensive rats leads to persistently reduced kidney renin and low blood pressure

**DOI:** 10.1101/2023.07.30.551193

**Authors:** Sean G. Byars, Priscilla Prestes, Vara Suphapimol, Fumihiko Takeuchi, Nathan De Vries, Michelle C Maier, Mariana Melo, David Balding, Nilesh Samani, Andrew Allen, Norihiro Kato, Jennifer L Wilkinson-Berka, Fadi Charchar, Stephen B Harrap

**Author notes:** **Name, email address, and complete address of corresponding author** Professor Emeritus Stephen Harrap, Department of Anatomy and Physiology University of Melbourne, VIC 3010, Australia.

## Abstract

**BACKGROUND:** Prevention of human hypertension is an important challenge and has been achieved in experimental models. Brief treatment with renin-angiotensin system (RAS) inhibitors permanently reduces the genetic hypertension of the spontaneously hypertensive rat (SHR). The kidney is involved in this reprogramming, but relevant genetic changes are unknown.

**METHODS:** In SHR, we studied the effect of treatment between 10 and 14 weeks of age with the angiotensin receptor blocker, losartan, or the angiotensin-converting enzyme (ACE) inhibitor, perindopril (with controls for non-specific effects of lowering BP) on differential RNA expression, DNA methylation and renin immunolabelling in the kidney at 20 weeks of age.

**RESULTS:** RNA sequencing revealed a 6-fold increase in renin gene (*Ren*) expression during losartan treatment (P < 0.0001). At 20 weeks, six weeks after treatment cessation, mean arterial pressure remained lower in the treated SHR (P = 0.006), kidney *Ren* expression was reduced by 23% (P = 0.03) and DNA methylation within the *Ren* promoter region was increased (P = 0.04). Experiments with the ACE inhibitor perindopril confirmed a long-term reduction in kidney *Ren* expression of 43% (P = 1.4 x 10^-6^). Renin immunolabelling was also lower after losartan or perindopril treatment (P = 0.002). RNA sequencing identified differential expression of 13 candidate genes (*Grhl1*, *Ammecr1l*, *Hs6st1*, *Nfil3*, *Fam221a*, *Lmo4*, *Adamts1*, *Cish*, *Hif3a*, *Bcl6*, *Rad54l2*, *Adap1*, *Dok4*) and the miRNA miR-145-3p. We found correlations between expression of mRNAs, miRNAs and lncRNAs that we believe represent genetic networks underpinning the decreased *Ren* expression and lower BP. Gene ontogeny analyses revealed that these networks were enriched with genes relevant to BP, RAS and the kidneys.

**CONCLUSIONS:** Early RAS inhibition in SHR reprograms genetic pathways and networks resulting in a legacy of reduced *Ren* expression and the persistent reduction in BP.

## INTRODUCTION

The ability to prevent the development of hypertension by treatment in early life would be an important advance in human health.^1^ This possibility has been raised by extensive research in the Spontaneously Hypertensive rat (SHR) model of human hypertension. Numerous independent studies have shown that administration of inhibitors of the renin-angiotensin system (RAS) such as angiotensin converting enzyme inhibitors (ACEi)^2, 3^ or angiotensin receptor blockers (ARB)^4^ in the prehypertensive period result in a permanent reduction in SHR blood pressure (BP) and increased lifespan.^5^

RAS-inhibiting drugs are common and effective treatments for established human hypertension, but their potential for preventing hypertension is less clear. Three human trials have attempted to emulate the SHR prevention paradigm, but with limited success.^1^ Effective prevention in humans is likely to depend on the appropriate age of treatment, length of treatment and selection of subjects. A more detailed understanding of the molecular basis of the legacy phenomenon in SHR should hold the key to effective prevention of hypertension in humans.

Several considerations are important here. First, the persistent effects following RAS inhibition in SHR are not observed following treatment with other agents such as vasodilators (e.g., hydralazine), diuretics, alpha-adrenergic blockers or calcium antagonists.^6–12^ This suggests the RAS is involved in both the genetic origins and prevention of SHR hypertension. Secondly, the legacy phenomenon is observed in some other hypertensive strains^13–16^ but not all,^17, 18^ suggesting a genetic predisposition to the legacy phenomenon.

In the current study, we tested our hypothesis that early RAS inhibition reprograms the genetically determined renal components of the SHR genetic hypertension. Our focus on the kidney is based on evidence in young SHR that abnormalities in renal hemodynamics and the RAS are important in the pathogenesis of the genetic hypertension. The prehypertensive SHR has reduced renal blood flow (RBF) and glomerular filtration rate (GFR) and high levels of kidney renin^19, 20^ and renal abnormalities have been linked to the inheritance of BP in cross-breeding experiments.^21^ The early renal hemodynamic abnormalities in SHR are reversed acutely by RAS inhibition^22^ and ameliorated in the long-term after early RAS inhibition.^2^ The importance of the kidney in SHR is emphasized by the results of renal transplantation between previously RAS-inhibited and untreated SHR in which BP “follows the kidneys” such that kidneys from treated SHR will lower BP in untreated animals and kidneys from untreated SHR will increase BP in the treated animals.^23^

We addressed the following specific questions in this study: 1) what are the long-term effects on the expression of RAS relevant genes following early RAS inhibition? 2) what other changes in coding and non-coding gene (miRNAs, lncRNAs) expression^24^ are associated with long-term effects? 3) is there evidence of differential DNA methylation that might explain changes in expression? 4) do differentially expressed genes show patterns of co-regulation? 5) are there identifiable clusters of differentially expressed genes relevant to biological processes, in particular BP? 6) is there evidence of a class effect for RAS inhibitors? and 7) are there genomic sequence variants in SHR that might predispose to long-term changes in RNA expression?

## METHODS

### Experimental outline

The basic experimental paradigm depicted in Figure 1 involved treatment of male SHR from 10 to 14 weeks of age with cardiovascular measurements and molecular and histological analyses of renal cortices at 14 and 20 weeks of age. All experiments were approved by The University of Melbourne Animal Ethics Committee (Ethics number 1313035) and were conducted in accordance with the Australian Code of Practice for the Care and Use of Animals for Scientific Purposes. We obtained 6-week-old inbred male SHR from the Animal Resources Centre (Canning Vale, Western Australia). All treatments were delivered via minipumps (Alzet model 2004, Durect Corp Cupertino, California).

**Figure 1:**
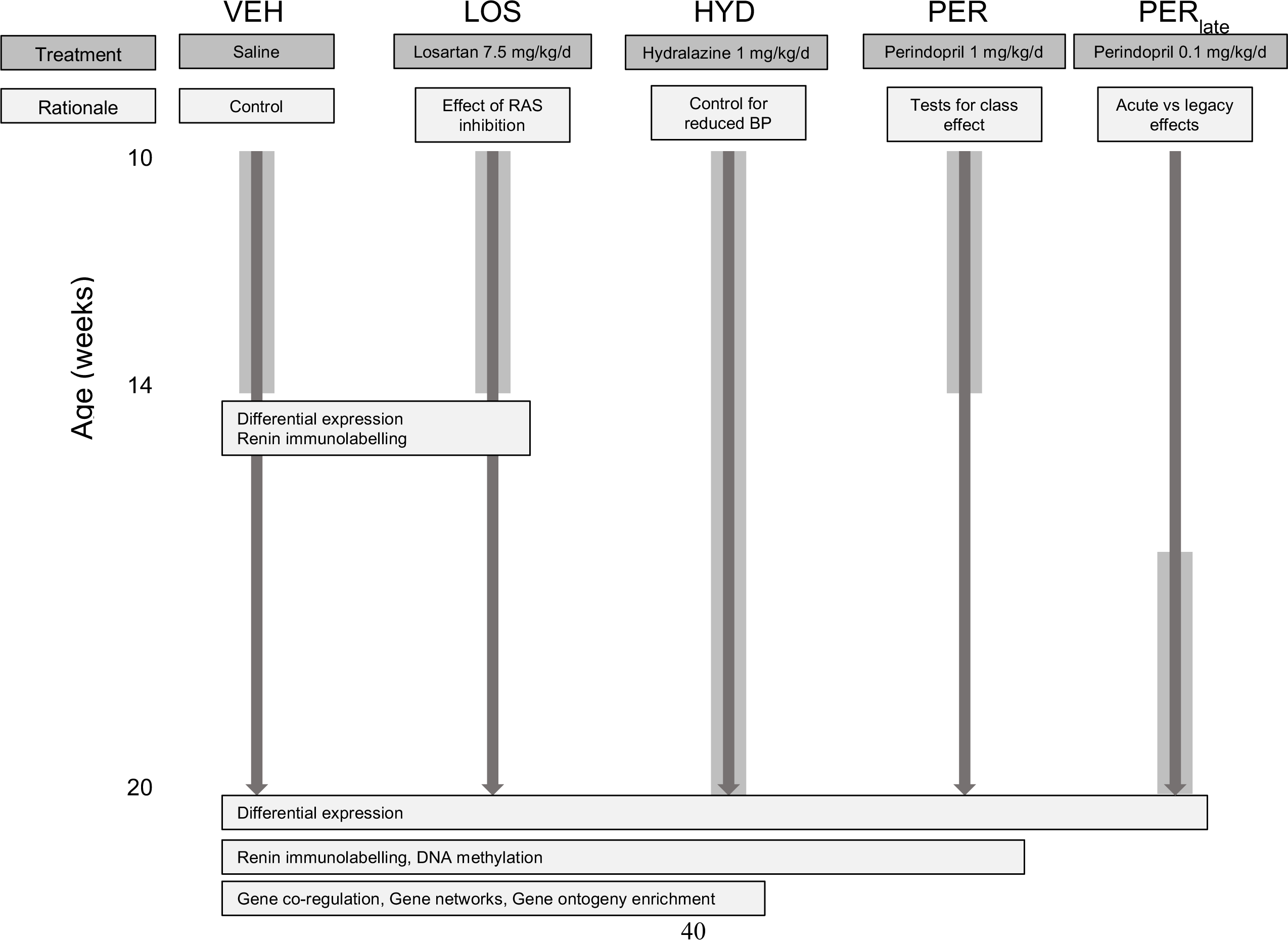
Outline of the basic experimental design showing the 5 experimental groups, their treatments and dosages and the experimental rationale for each group. The treatment periods are shown by the wide grey vertical bars against age in weeks. The data obtained are shown in boxes at 14 and 20 weeks of age. The boxes span the groups for which data was analysed.

#### 1. Losartan experiments

A primary set of experiments was based on a group of SHR treated with losartan from 10-14 weeks of age (LOS, 7.5 mg/kg/d, n = 18). Two control groups were used for identifying differential RNA expression. Our initial comparisons were made with vehicle (saline) treated SHR from 10-14 weeks of age (VEH, n =18) to detect overall differences resulting from losartan treatment. We utilised a second control group to account for differences in LOS that might be the result of non-specific effects BP reduction *per se*. For this purpose, we included a group of SHR that were actively treated with the vasodilator hydralazine from 10-20 weeks of age (HYD, 1.0 mg/kg/d, n = 6) to mimic the BP effects following RAS inhibition (Figure 1). Differential expression analyses focussed specifically on genes associated with the RAS and genome-wide analyses to identify candidate RNA species most strongly associated with the long-term effects following losartan at 20 weeks. These experiments provided detailed information regarding differential total mRNA and specific miRNA expression, DNA methylation and the presence and nature of gene co-regulation, gene networks and gene ontology enrichment.

#### 2. Perindopril experiments

We undertook additional experiments with the ACE inhibitor perindopril from 10-14 weeks of age (PER, 1.0 mg/kg/d, n = 11) to test specific hypotheses generated in the losartan experiments to seek evidence of a class effect of early RAS inhibition. Finally, to compare effects of active RAS inhibition versus the legacy of early treatment we included a group of SHR (PER_late_) that was treated with a lower dose of perindopril (0.1 mg/kg/d, n = 4) from 16-20 weeks of age to achieve a BP approximating the legacy effect following early RAS inhibition.

### BP measurement and relative cardiac mass

Arterial pressures were measured using radiotelemetry between 10 and 20 weeks of age in a sample from 4 groups (VEH n = 2, LOS n = 2, HYD n = 2, PER = 3). These animals were excluded from sequencing analyses in case of confounding effects of surgery and *in situ* telemetry probes. Telemeters were inserted into the abdominal aorta at age 7 weeks (see Detailed Methods). Between 10 and 20 weeks of age, recordings were made over one 24-h period every week, from which we obtained the average daylight, resting 12-h measurement of mean arterial pressure (MAP) for each rat for that week. We also measured tail cuff systolic blood pressure (SBP) in larger groups of animals. Animals were accustomed to the light restraint and BP measurement protocol and SBP was measured using a programmed sphygmomanometer with a pneumatic pulse transducer (PE-300; Narco Bio-System, Houston, TX). In addition, we used relative cardiac mass (RCM) as an indication of the average BP. At the end of experiments hearts were removed, blotted dry, weighed and RCM was calculated as the ratio of heart to body weight.

### Kidney cortex collection

Animals were rapidly euthanized using isoflurane and ketamine and one kidney from each animal was dissected on ice to obtain cortical tissue for RNA extraction as described in the Detailed Methods. The other kidney was immediately fixed in formalin for histological studies.

### RNA-sequencing and methylation analyses

Sequencing was performed on individual kidney cortex samples. Total RNA sequencing to capture coding and non-coding genes (eg mRNA, lncRNA, snoRNA), miRNA and methylation sequencing were performed by the Australian Genome Research Facility (AGRF, Melbourne, Australia). Quality control of total RNA-sequencing data and miRNA sequencing data with FastQC revealed high quality sequence and base scores (see Detailed Methods). Total RNA sequencing was also obtained from Novogene (Beijing, China) to permit confirmation of VEH and LOS differential expression results. A total of three sequencing runs by AGRF and Novogene were made to accommodate all samples and differential expression analyses were made only by comparison within individual runs to avoid potential batch effects. Differences in expression are presented as log base 2 fold changes (log_2_FC).

All total RNA sequencing utilized the same parameters including rRNA removal (Ribo-Zero depletion) and Illumina HiSeq 150bp paired-end sequencing at high depth (∼100 million reads per sample). Illumina NovaSeq (50bp single-end reads, ∼10-20 million reads per sample) was used for miRNA sequencing. Illumina HiSeq (100bp single-end reads, 30 million reads per sample) was used for methylation sequencing using the reduced representation bisulfite sequencing (RRBS) technique. For all sequencing types, details regarding the assessment of read quality, alignment and quantification, data normalisation and analyses of differential expression are provided in the Detailed Methods.

### Gene co-regulation, gene network and gene ontogeny enrichment analyses

To explore potential gene co-regulation, gene networks and enriched gene ontology terms related to LOS treatment, we undertook a complimentary series of analyses, the details for which are provided in the Detailed Methods.

First, we sought to investigate co-regulatory relationships between differentially expressed protein-coding and non-coding RNA species to help explore possible changes in the regulatory machinery following losartan treatment. We employed expression data from the total RNA (Table S1) and miRNA (Table S2) data sets in multivariate analyses using regularized Canonical Correlation Analysis (rCCA) and other tools in the R package mixOmics version 6.6.2.^25^

Next, we used Weighted Correlation Network Analysis (WGCNA version 1.68^26^) to identify networks comprising distinct clusters of co-expressed genes (modules) related to the long-term effects of LOS treatment. Three WGCNA networks were generated, one for each class of studied genes (mRNA, small-RNAs and miRNA). Expression data from genes related to LOS treatment was used to build these networks (Tables S3-S5) and selected so that leading differentially expressed genes were prioritised while also including sufficient numbers to allow network construction (see Detailed Methods). Within each gene module, hub genes were identified, as these may be particularly important to consider related to LOS treatment given their central roles in gene networks.^25^ Finally, we correlated module eigengenes (ME) with BP to examine gene by trait correlations relevant to LOS treatment effects. An ME represents each module’s summary expression profile and is a useful tool for studying relationships between different modules and relevant physiological phenotypes.

To obtain insights into biological pathways that may be altered in response to treatment, we undertook gene ontology enrichment analyses (see Detailed Methods) using leading differentially expressed genes in the LOS experiments.

### Renin immunolabelling

Immunohistochemistry for renin was performed as described previously^27, 28^ and all analyses were blinded to the treatment groups. Briefly, rat kidneys were fixed in 10% buffered formalin. Five μm paraffin sections of kidney were incubated with a polyclonal mouse renin protein antibody raised against pure mouse submandibular gland renin (1:8000). A negative control without the primary antibody was included (see Detailed Methods). For quantitation, four randomly chosen sections from each kidney at least 125 μm apart, were selected.

Images of renin immunolabelling associated with the juxtaglomerular apparatus (JGA) from 50 glomeruli per section were captured and the proportion of renin-positive JGAs per section were averaged across four sections from each sample.

### Renin gene sequence variants

We searched for DNA variants in SHR-derived strains (SHR/SHRSP) that might predispose to the long-term antihypertensive effects following RAS inhibition. We used whole genome sequencing data variants in and around the renin gene for SHR/Utx: a substrain of University of Texas, Houston, SHRSP/Bbb and WKY/Bbb: substrains of Max-Delbruck Center for Molecular Medicine;^29^ SHRSP/Izm, SHR/Izm and WKY/Izm: substrains of Japanese colony.^30^ We aligned the NGS reads to the BN reference (mRatBN7.2) for practical reasons in the interval of 44,781,715-44,808,987 on RNO13, and sought DNA variation (SNPs, insertions, deletions, nucleotide repeats) in SHR and SHRSP that were not observed in WKY or BN sequences. Transcription factor (TF) binding sites were predicted using PROMO (https://alggen.lsi.upc.es/cgi-bin/promo_v3/promo/promoinit.cgi?dirDB=TF_8.3) with humans selected as factor’s species, considering the reported similarities of TF motifs between mammals.^31^

### Statistical analyses

Parametric analyses (t-tests, ANOVA, repeated measures ANOVA) were used to compare data that are close to normally distributed and presented as mean ± standard deviation (SD) unless otherwise specified. Non-parametric tests (Mann-Whitney test, Kruskal-Wallis test) were used for non-normal data that are presented as median and interquartile range (IQR) unless otherwise stated. Tests were performed using IBM SPSS Statistics version 27.

Specialised statistical approaches are described in the relevant sections in the Detailed Methods.

## RESULTS

### Losartan experiments

#### Cardiovascular effects of treatment

The direct telemetric MAP recordings (Figure 2) demonstrated that losartan (P = 0.005 by repeated measures ANOVA) and hydralazine (P = 0.009) significantly reduced BP during treatment from 10-14 weeks of age. On cessation of treatment, the BP of losartan-treated SHR rose to levels similar to the SHR receiving ongoing hydralazine treatment and both remained significantly lower than control SHR (LOS P = 0.006; HYD P = 0.005). The tail cuff SBP recordings (mean ± SD) confirmed these differences at 14 weeks (VEH: n = 7, 206±16 mmHg; LOS: n = 10, 145±22 mmHg; HYD: n = 5, 153±12 mmHg) (ANOVA P < 0.0001) and 20 weeks (VEH: n = 8, 220±18 mmHg; LOS: n = 8, 187±16 mmHg; HYD: n = 3, 153±8 mmHg) (ANOVA P < 0.0001). At 14 weeks of age mean RCM was significantly reduced in LOS (n = 6, 3.21±0.14 g/kg) and HYD (n = 2, 3.46±0.01 g/kg) compared with the VEH (n = 4, 3.63±0.13 g/kg) (ANOVA P = 0.002) group. Similarly, at 20 weeks of age RCM was significantly less in LOS (n = 11, 3.36±0.12 g/kg) and HYD (n = 3, 3.34±0.28 g/kg) compared with VEH (n = 8, 3.61±0.12 g/kg) (ANOVA P = 0.001).

**Figure 2:**
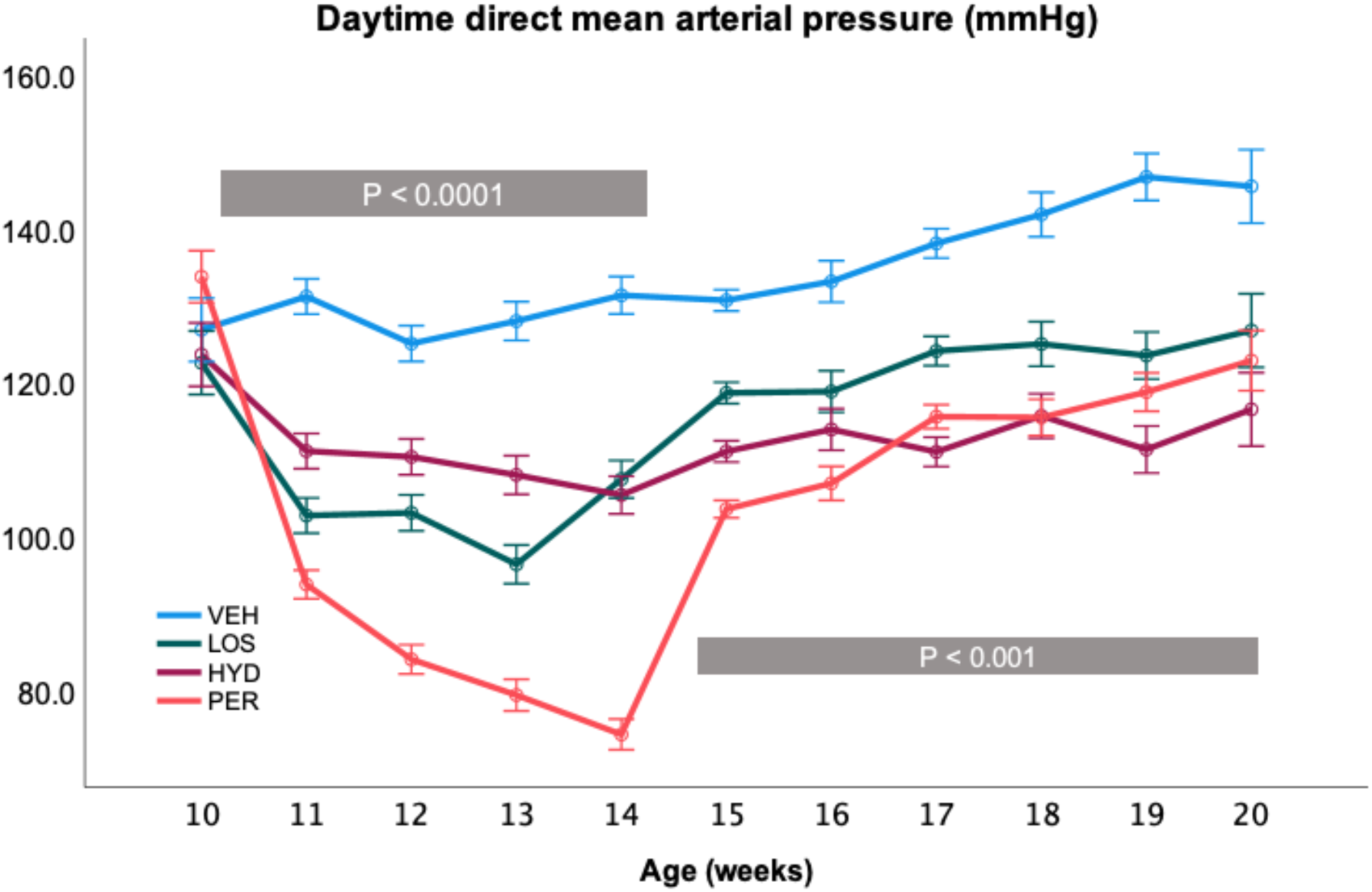
Direct mean arterial pressure measured in male SHR from 10 to 20 weeks of age. Animals received saline (VEH, n = 2, blue), losartan (LOS, n = 2, green) or perindopril (PER, n = 3, orange) between 10 and 14 weeks of age or hydralazine (HYD, n = 2, purple) between 10 and 20 weeks of age. Weekly values are summarised as estimated marginal means from the repeated measures ANOVA and error bars represent the SEM. The P values represent the overall effect of treatment during (10 to 14 weeks) and after treatment for LOS and PER (15 to 20 weeks) by repeated measures ANOVA. Individual treatment comparisons with VEH are given in the text.

#### RAS gene expression

During losartan treatment at 14 weeks there were, as anticipated, significant changes in the expression of several genes involved in the RAS and its signalling pathways (Table S6). The most prominent was a significant increase in renin gene (*Ren*) mRNA (log_2_FC = 2.46, FDR q = 1.3 x 10^-133^). Other significant changes included increased expression of the genes encoding angiotensinogen (*Agt*: log_2_FC = 0.50, FDR q = 3.2 x 10^-5^), angiotensin II receptor associated protein^32^ (*Agtrap*: log_2_FC = 0.27, FDR q = 2.2 x 10^-5^) and transforming growth factor beta 1^33^ (*Tgfb1*: log_2_FC = 0.38, FDR q = 2.1 x 10^-5^) and reduced expression of renin receptor ATPase H+ transporting accessory protein 2^34^ (*Atp6ap2*: log_2_FC = -0.22, FDR q = 4.3 x 10^-5^), mitogen-activated protein kinase 1^35^ (*Mapk1*: log_2_FC = -0.25, FDR q = 0.004) and SRY-box transcription factor 6^36^ (*Sox6*: log_2_FC = -0.25, FDR q = 0.018).

At 20 weeks of age, *Ren* was the only RAS gene to show significant differential expression following prior losartan treatment, with expression reduced compared with both VEH (log_2_FC = -0.31, P = 9.5 x 10^-6^, Table S7) and HYD (log_2_FC = -0.37, P = 0.03, Table S8).

#### Genome-wide differential expression analyses

Comparing the renal cortices from 20-week-old LOS and VEH SHR, we identified 164 differentially expressed genes (Table S7) sequenced by AGRF and 177 differentially expressed genes in the Novogene sequencing results from the same samples (Table S9). There were 35 genes that showed significant (FDR q < 0.05) and consistent (in terms of direction and magnitude) differential expression in both the AGRF and Novogene analyses (Figure S1) and we designated these genes as “validated” (Table 1).

**Table 1:**
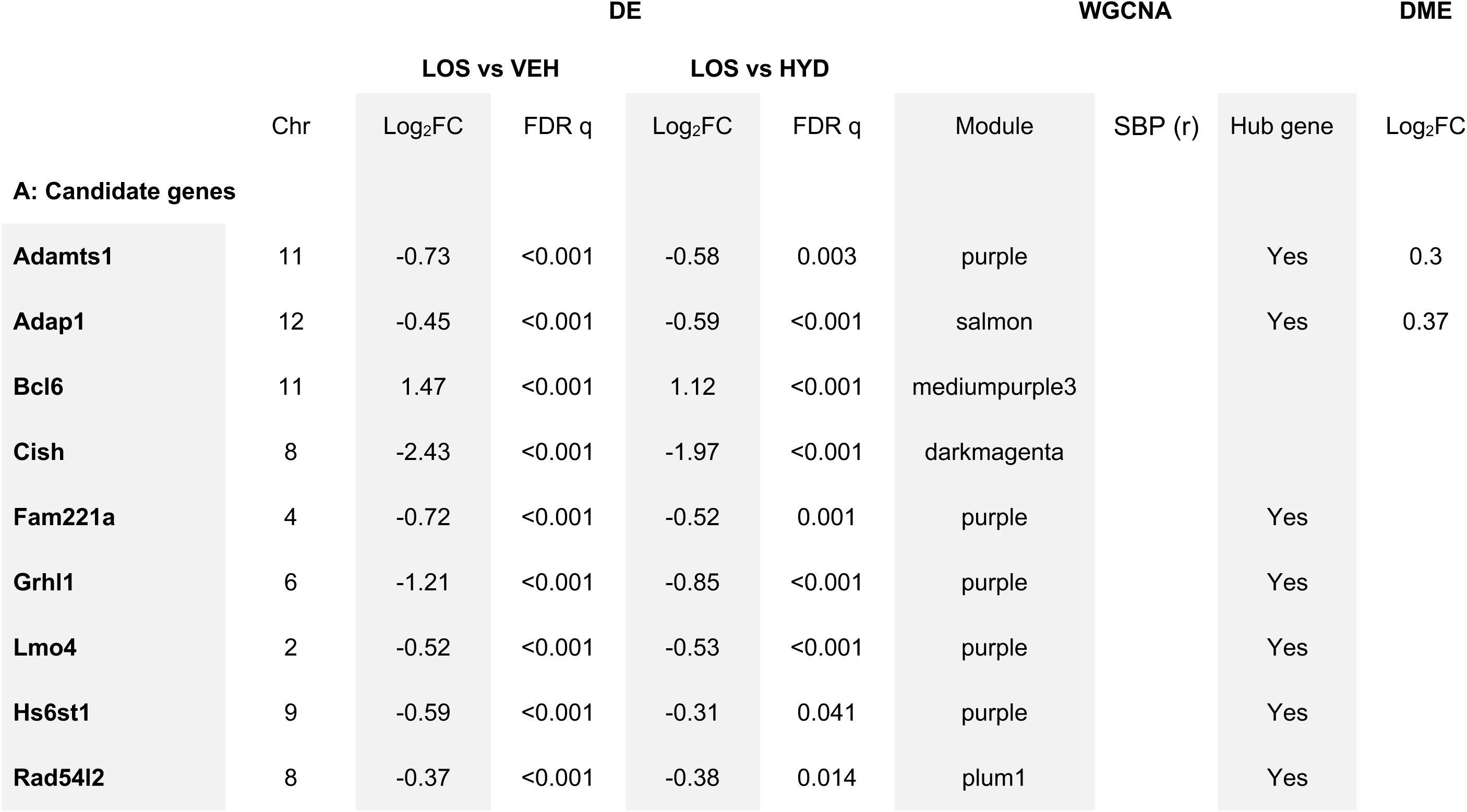
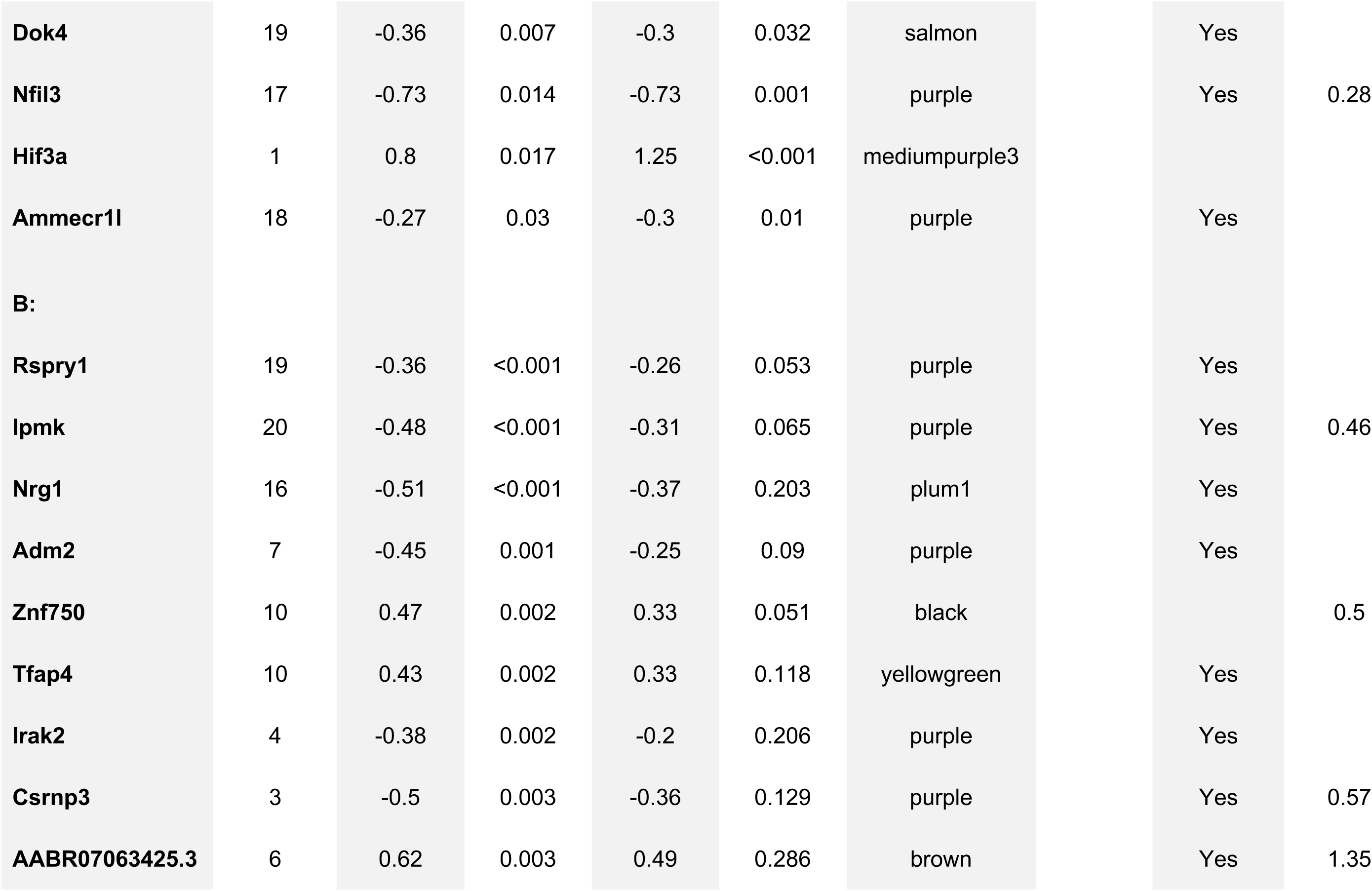
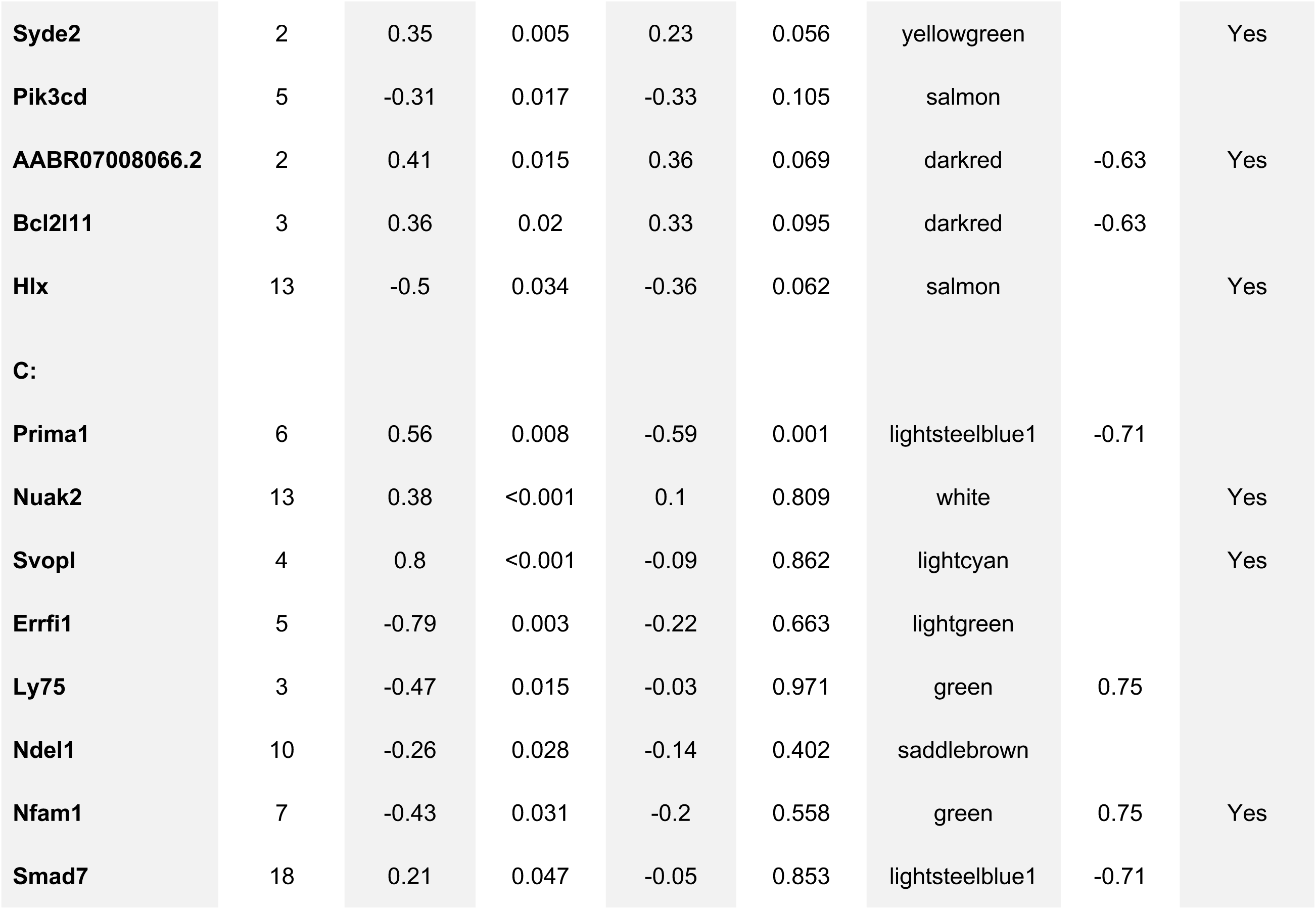
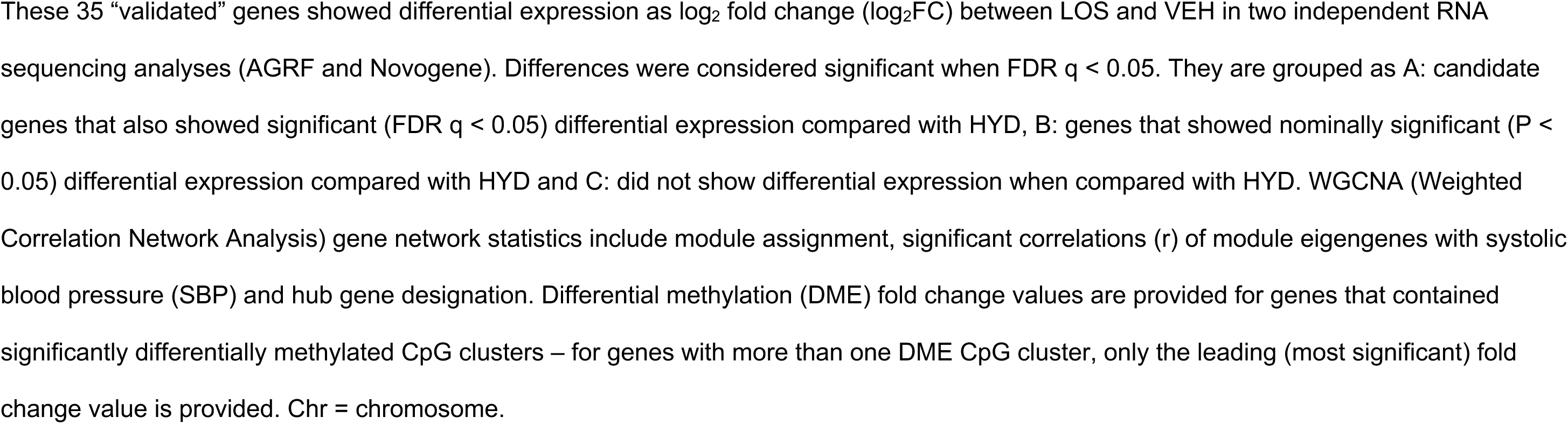
Genes showing differential expression (DE) in kidney cortices at 20 weeks of age after prior treatment with losartan.

Using HYD as controls (Table S8), we identified 13 genes among the 35 validated genes that showed significant differential expression in the same direction as VEH analyses when compared with HYD (Table 1A). As their expression could not be ascribed to lower BP *per se*, we nominated these 13 loci as “candidates” for the long-term effects of losartan: *Grhl1*, *Ammecr1l*, *Hs6st1*, *Nfil3*, *Fam221a*, *Lmo4*, *Adamts1*, *Cish*, *Hif3a*, *Bcl6*, *Rad54l2*, *Adap1*, *Dok4* (see Table 1A). Another 14 genes showed significant differential expression against VEH but only nominal differences with HYD (Table 1B).

The expression patterns of these 13 candidates at 14 and 20 weeks are shown in Figure 3. Although all candidates showed significant differential expression at 20 weeks, only 3 candidates: *Nfil3* (log_2_FC = -1.54, FDR q = 6.6 x 10^-13^), *Adap1* (log_2_FC = 0.53, FDR q = 2.8 x 10^-6^) and *Dok4* (log_2_FC = 0.25, FDR q = 0.007), were differentially expressed during active losartan treatment at 14 weeks; suggesting they are sensitive to the effects of losartan itself.

**Figure 3:**
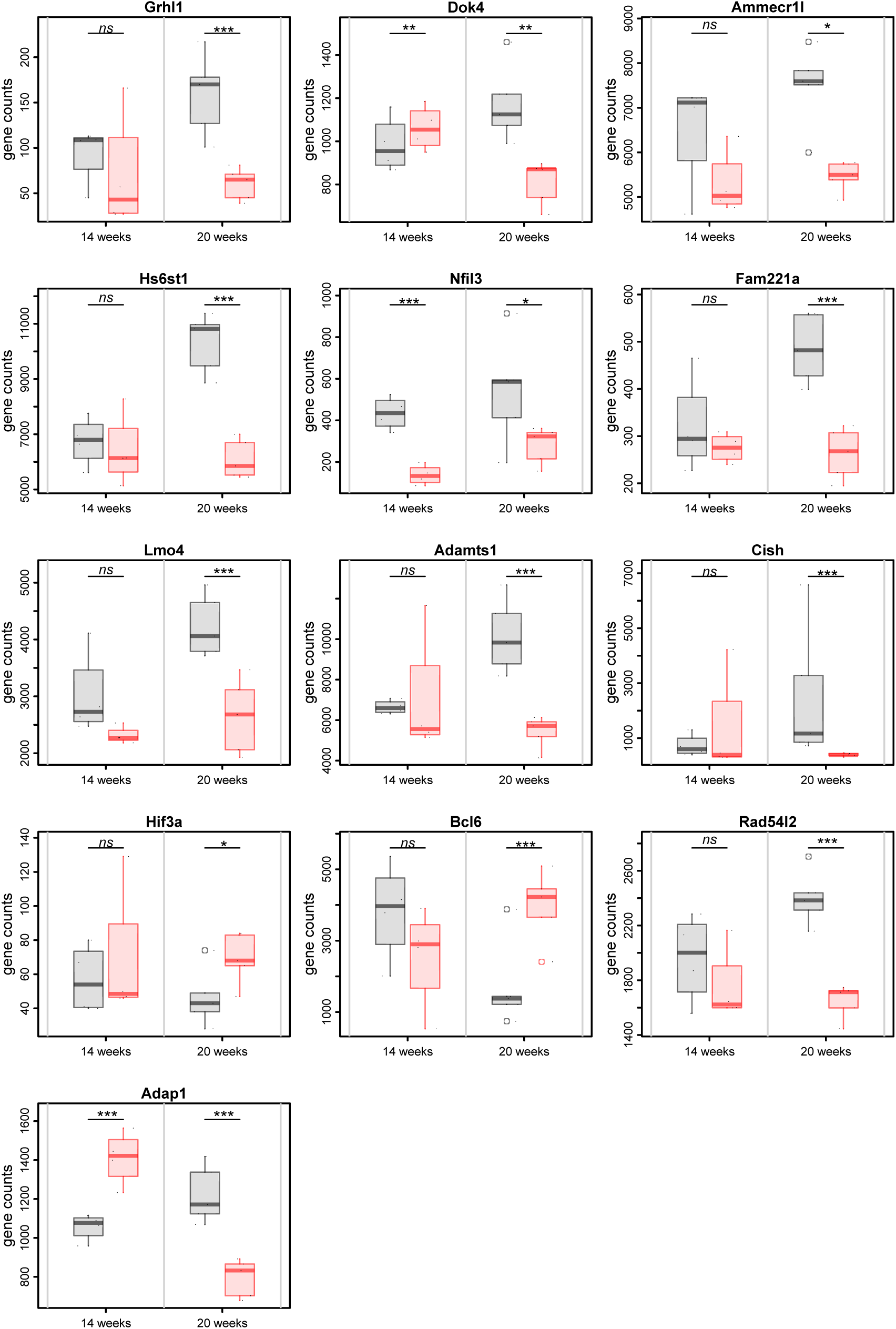
Changes in expression of 13 candidate (of the 35 validated) genes at 14 and 20 weeks in losartan and control SHR animals. Gene expression (counts) are displayed individually (black points) and summarized as boxplots (vehicle: grey, losartan: red). FDR q-values (***<0.001; **<0.01; *<0.05; ns>0.05) are from differential gene expression analyses.

In addition, we identified 45 miRNAs that showed differential expression between the LOS and VEH groups at 20 weeks (Table 2). Most of these miRNAs (n = 33) showed up-regulation in the LOS animals compared with controls.

**Table 2.**
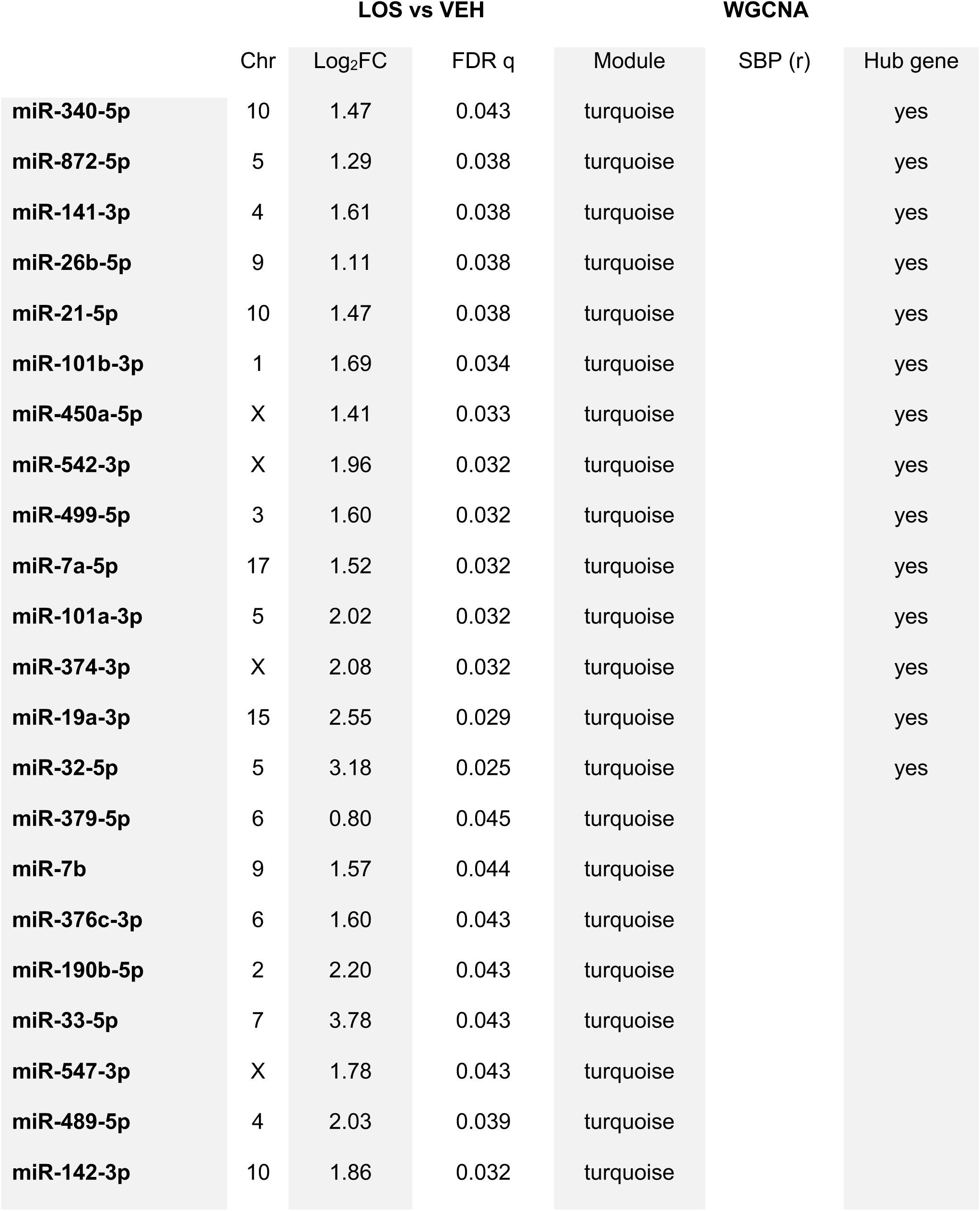
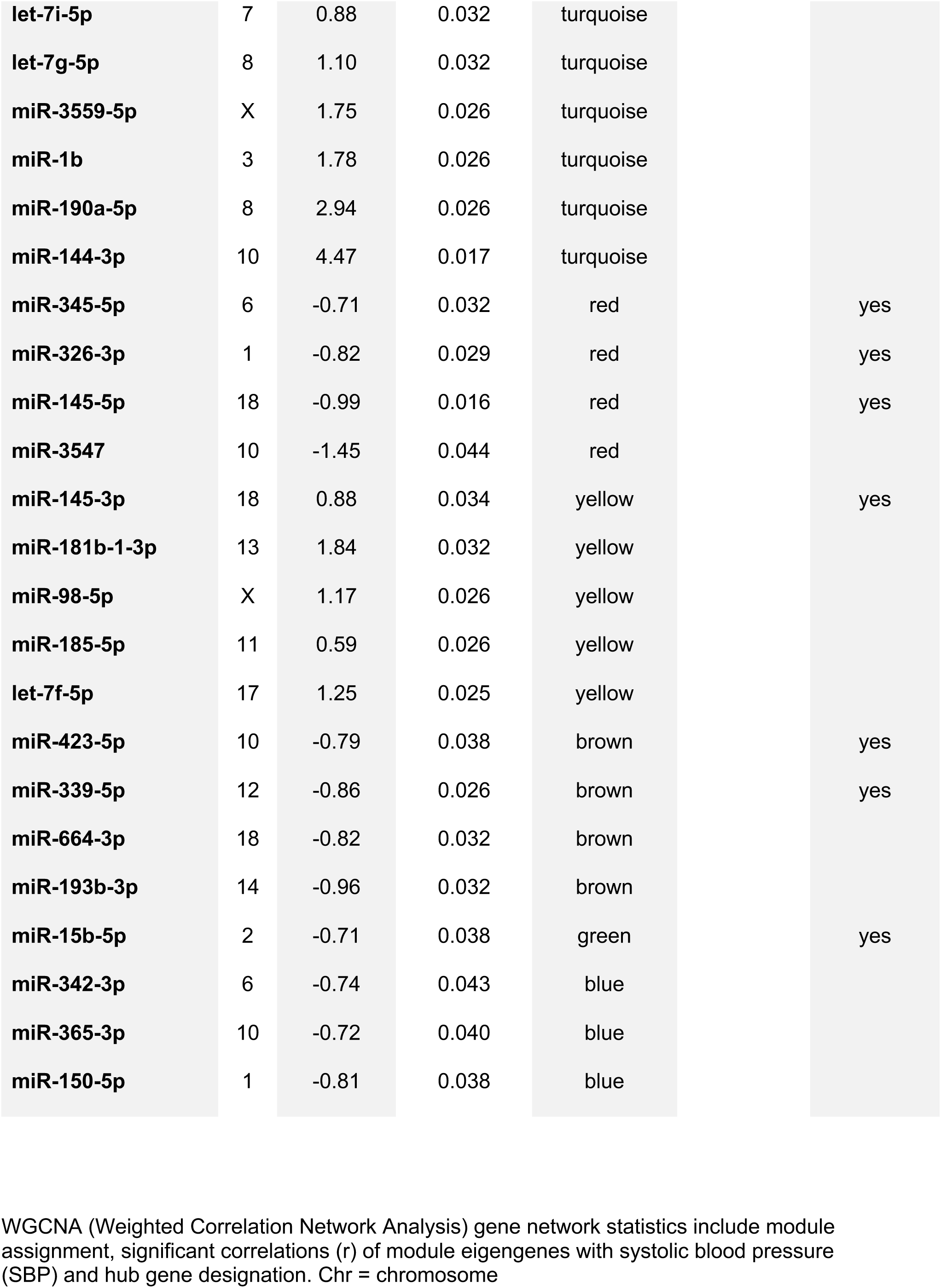
miRNAs showing differential expression at 20 weeks in response to losartan treatment in the spontaneously hypertensive rat.

#### Correlations between coding and non-coding RNA

Given the potential control of gene expression by non-coding RNA, we investigated multivariate correlations between the differentially expressed coding and non-coding RNAs. Those which are highly correlated (|r| ≥ 0.8) are shown in Figure S2. When considering large negative correlations (r < -0.8) between the 13 candidate genes and non-coding RNAs (Figure 4, Table S10), we found that the miRNA miR-145-3p was highly negatively correlated with nine of the 13 candidates: *Adamts1*, *Hs6st1*, *Fam221a*, *Grhl1*, *Ammecr1l*, *Adap1*, *Dok4*, *Lmo4*, *Rad54l2* (Table S10, Figure S2C). We also noted that among the lncRNAs, AC115371.^37^ was very highly correlated with numerous miRNAs (n = 29 of 45) and showed the highest correlation (r = 0.97) with miR-145-3p (Table S11, Figure S3).

**Figure 4.**
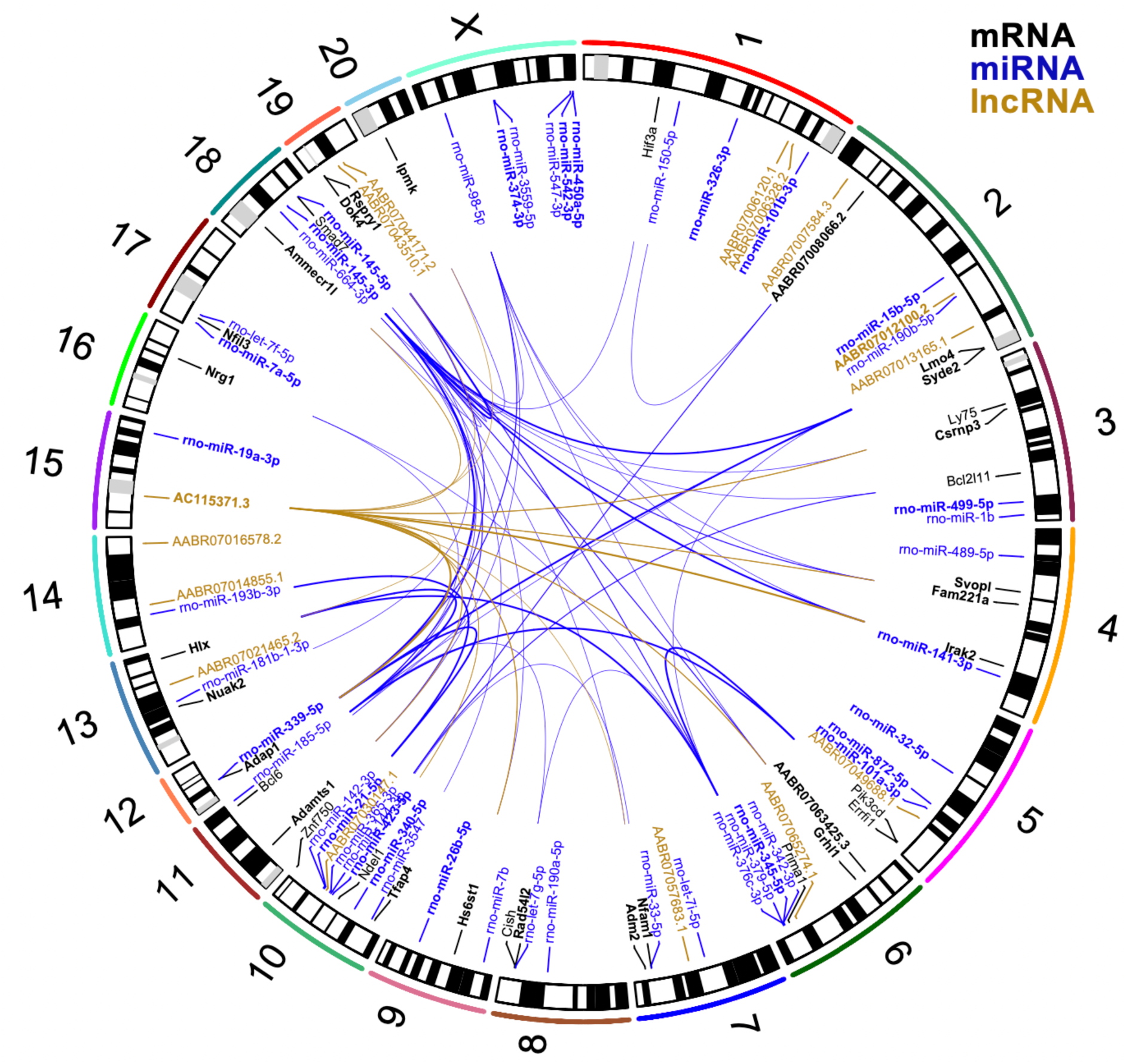
Leading differentially expressed genes and their associations. Including 35 mRNAs (black text) from Table 1, 45 miRNAs (blue text) from Table 2 and 15 lncRNAs (gold text). Genes are in bold text if they were identified as hub genes from the WGCNA analysis. For each of the 35 mRNAs, the leading two negative correlations < -0.8 (if present) with non-coding genes are presented in the centre, with thicker lines representing larger negative associations (for a full set of multivariate correlations, see Figure S2 and Table S10).

#### Gene network analyses, modules and SBP correlations

The WGCNA network analyses identified a total of 43 modules among 2844 protein coding genes (Table S3, Figure S4A) ranging in size from 25 to 207 genes each. Among the 13 candidate genes, ten were identified as hub genes (Table 1A) suggesting a central role within these gene co-expression modules related to the legacy of losartan treatment. Seven of these (*Grhl1*, *Ammecr1l*, *Hs6st1*, *Nfil3*, *Fam221a*, *Lmo4*, *Adamts1*) were in the module purple (Table S3). The genes *Adap1* and *Dok4* were in module salmon and interestingly these two genes had shown increased expression during losartan treatment at 14 weeks but significantly reduced expression at 20 weeks. The non-coding RNA species from the total RNA sequencing (n = 186) were allocated separately to 19 modules (Table S4, Figure S4B), with miRNAs, snoRNAs and lncRNAs (including AC115371.3) identified as hub genes. The miRNAs (n = 210) were grouped into seven modules (Table S5, Figure S4C).

The correlation between SBP and the summary expression (eigengene) for each module was examined and found to be significantly correlated with nine mRNA modules, three non-coding RNA modules and one miRNA module (Figure S4).

#### Gene ontogeny enrichment analysis

Differentially expressed genes related to 20-week LOS treatment showed significant enrichment (FDR q < 0.05) with 153 biological processes (Table S12B). Given that many enrichment terms were very broad (i.e., including thousands of genes), we also explored nominally significant (P < 0.05) terms. We identified ten terms that involved regulation of BP and/or RAS (Table S12A) that were closely related (Table S12D). Within these domains there were two differentially expressed RAS-related genes (*Ren*, *Agt*) in nominally significant RAS-related pathways such as ‘regulation of blood volume by renin-angiotensin’ (GO:0002016, P = 0.002, Table S12C). There were also other differentially expressed genes (e.g., *Adm2*, *Bdkrb2*, *Edn3*, *Pde4d*) found in nominally significant parent terms such as ‘positive regulation of BP’ (GO:0045777, P = 0.003).

#### Co-regulation of RAS gene expression

To explore potential co-regulation between RAS-specific and other differentially expressed genes in LOS animals, we investigated the multivariate correlation between *Ren* (and other relevant RAS genes) and the validated differentially expressed 35 mRNA and 45 miRNA at 20 weeks and with the 35 mRNA at 14 weeks during treatment (Tables S13-S15, Figure S5, Detailed methods).

**Figure 5:**
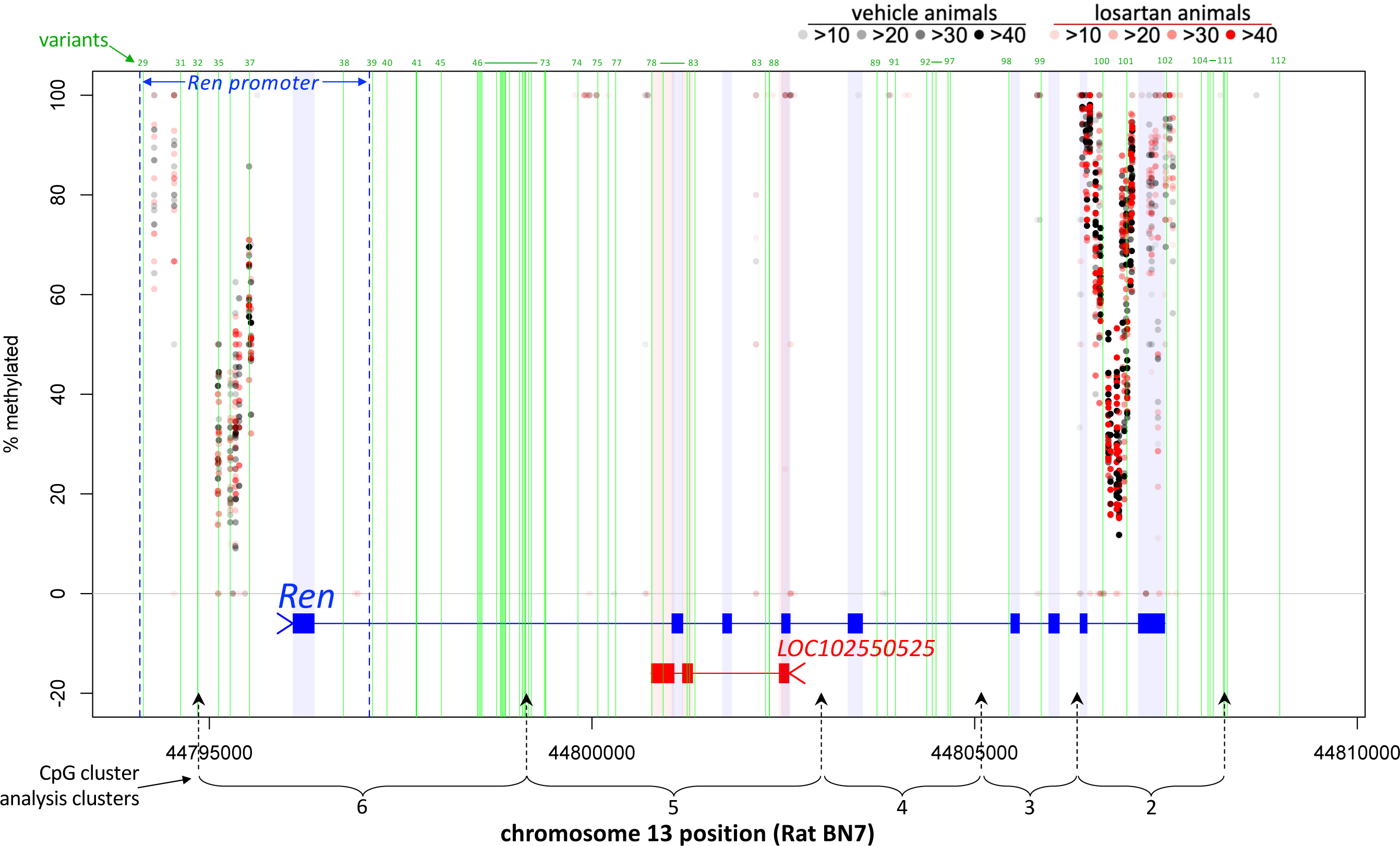
Patterns of methylation across *Ren* and LOC102550525. Detailed view of methylation % by vehicle and losartan animals (point shading reflects count number per CpG). Green vertical lines show position of variants (see variant description in Table S18) in and around *Ren*. General promoter region for *Ren* indicated by dark blue dashed lines. Position of CpG clusters (2-6) across *Ren* shown which corresponds to cluster number in CpG cluster analysis (Table S16).

Among the RAS genes, *Ren* was most frequently correlated (n = 18 genes with |r| ≥ 0.8) with the 35 validated genes at 20 weeks (Table S13, Figure S5A). This included six out of the 13 candidate genes, including the two strongest correlations (with *Bcl6* and *Hs6st1*). Only one other RAS gene (*Agt*) was highly correlated with three of the 35 validated genes at 20 weeks (Table S13).

The opposite was apparent at 14 weeks (Figure S5B). *Ren* (despite having markedly elevated expression) was less highly correlated (n = 6 genes with |r| ≥ 0.8) with the 35 validated genes, and only three of these (e.g., *Nfil3*, *Dok4* and *Adap1*) were among the 13 candidate genes (Table S14). Many of the other RAS genes (*Agt*, *Agtrap*, *Atp6ap2*, *Mapk1*) that were not highly correlated at 20 weeks, however, were so at 14 weeks.

In terms of correlations between miRNA and RAS genes at 20 weeks of age (Table S15) the highest correlation observed was between *Ren* and the miRNA miR-664-3p (Figure S5C).

Figure S3 shows the observed strong intercorrelations between *Ren*, miRNA and lncRNA.

#### Differential methylation

Among the 13 candidate genes, the cluster-based analysis revealed significantly increased methylation at CpG clusters in 20-week LOS SHR for *Adamts1*, *Adap1* and *Nfil3* (Table 1A, Table S16) corresponding with their lower RNA expression. Among the RAS genes only *Ren* showed evidence of increased methylation in cluster 2 (Table S16) that was localised in the 3’ end of *Ren* as shown in Figure 5. In analyses focused on gene promoter regions (Table S17) among the 35 validated and RAS genes, only *Ren* showed significantly increased methylation (Figure 5, log_2_FC = 0.30, permutation p-value 0.04) at 20 weeks.

### Perindopril experiments

#### Cardiovascular effects of treatment

The direct MAP recordings in PER SHR are show in Figure 2 with significant reductions in pressure during (P = 0.0003 by repeated measures ANOVA) and after (P = 0.004) treatment. Compared with VEH, mean tail cuff SBP in the PER group was significantly reduced during (PER: n = 4, 125±15 mmHg vs VEH: n = 4, 215±10 mmHg, t-test P < 0.0001) and after (PER: n = 10, 161±17 mmHg vs VEH: n = 8, 220±18 mmHg, t-test P < 0.0001) treatment. This pattern was mirrored by the lower RCM at 14 (PER: n = 4, 2.81±0.09 g/kg vs VEH: n = 4, 3.63±0.13 g/kg, t-test P = 0.0001) and 20 (PER: n = 10, 3.27±0.15 g/kg vs VEH: n = 12, 3.65±0.26 g/kg, t-test P = 0.0001) weeks of age.

#### Kidney Ren expression

We found reduced kidney cortex *Ren* expression compared with HYD at 20 weeks PER (log_2_FC: -0.80, P = 1.4 x 10^-6^). Active treatment with perindopril (0.1 mg/kg/d) between 16 and 20 weeks of age resulted in reduced SBP (n = 4, 179±12 mmHg, P = 0.002) and RCM (3.24±0.16 g/kg, P = 0.001) similar to PER animals. However, in distinct contrast to the reduction in *Ren* expression in PER animals, the PER_late_ animals showed was a 3-fold increase in *Ren* expression (log_2_FC: 1.57, P = 7.6 x 10^-49^).

#### Candidate gene expression

Although among the 13 candidates identified in the losartan experiments, 9 showed differential expression after perindopril in the same direction as losartan, only one showed significant change in the perindopril experiments – *Nfil3*. *Nfil3* expression was reduced in the kidneys of PER (log_2_FC -1.09, FDR q = 1.4 x 10^-16^). In addition, active perindopril treatment at 20 weeks was also associated with a significant reduction in *Nifl3* expression (log_2_FC - 0.79, P = 1.2 x 10^-8^, FDR q = 2.2 x 10^-6^), mirroring what had been observed at 14 weeks with losartan treatment.

### Renin immunolabelling

To complement the kidney *Ren* expression, we measured renin immunolabelling in the cortex of 14- and 20-week-old SHR (Figure 6). During treatment at 14 weeks, renin immunolabelling (summarised as median and interquartile range – IQR) was significantly higher in LOS SHR (n = 6, 45.2%, IQR 3.9%) compared with VEH (n = 4, 10.1%, IQR 6.5%) (P = 0.01 by Mann-Whitney test). However, at 20 weeks renin immunolabelling was lower in LOS (n = 7, 7.1%, IQR 1.3%) and PER (n = 8, 10.8%, IQR 7.3%) kidneys compared with VEH (n = 8, 13.7%, IQR 5.8%), while renal renin was higher in HYD kidneys (n = 6, 18.3%, IQR 2.37%). Renin immunolabelling after any RAS inhibition (LOS and PER) (7.6%, IQR 5.1%) was lower than VEH (P = 0.05 by Mann-Whitney test) and HYD (P = 0.001 by Mann-Whitney test). These changes corroborated the *Ren* RNA expression results.

**Figure 6.**
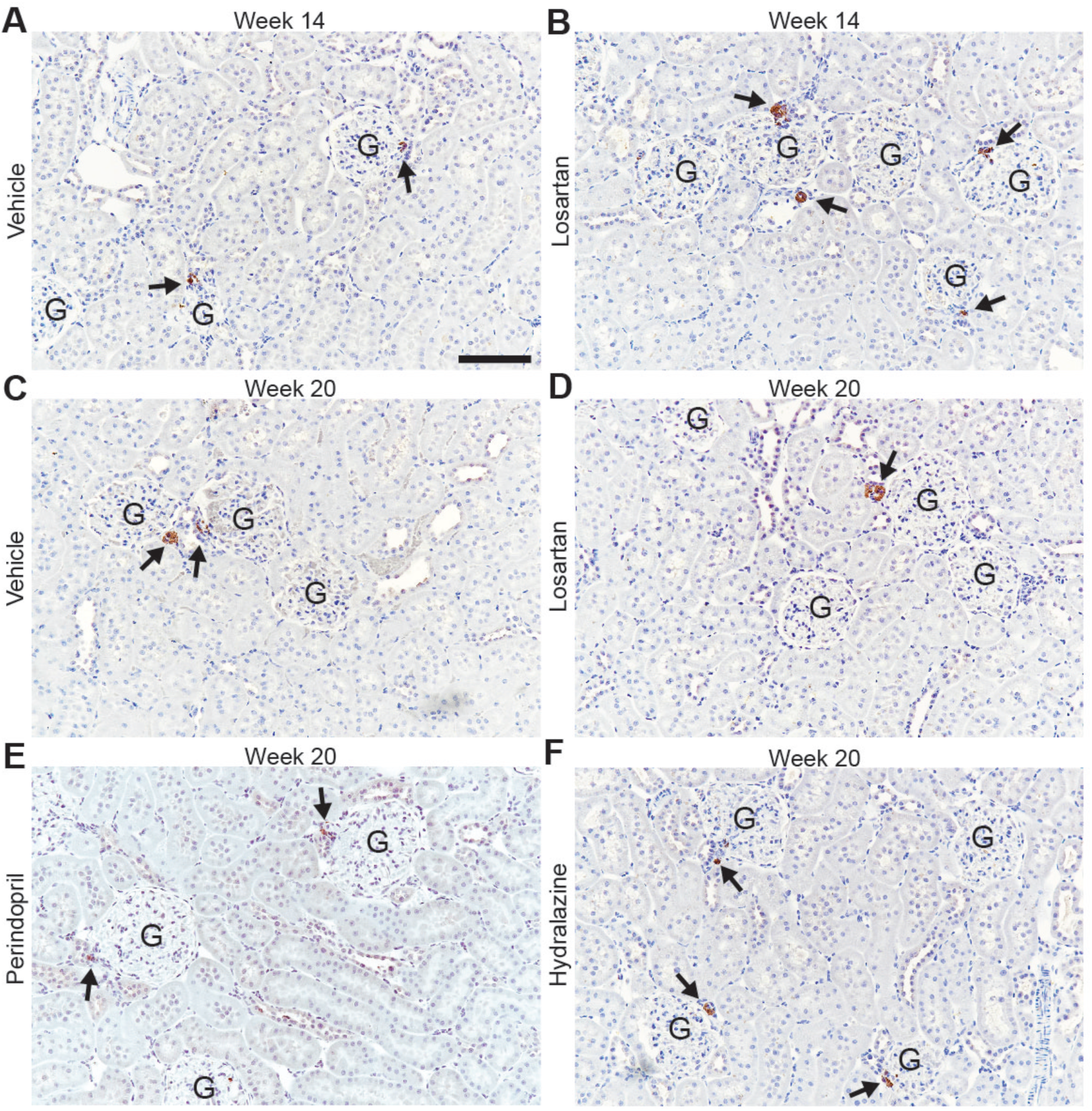
Renin immunolabelling in kidneys from 14-week (Panels A & B) and 20-week-old (Panels C – F) SHR treated with vehicle, losartan, perindopril and hydralazine. Three μm paraffin sections counterstained with hematoxylin. G, glomerulus. Arrows denote renin immunolabelling (brown). Scale bar, 100 μm.

**Figure 7:**
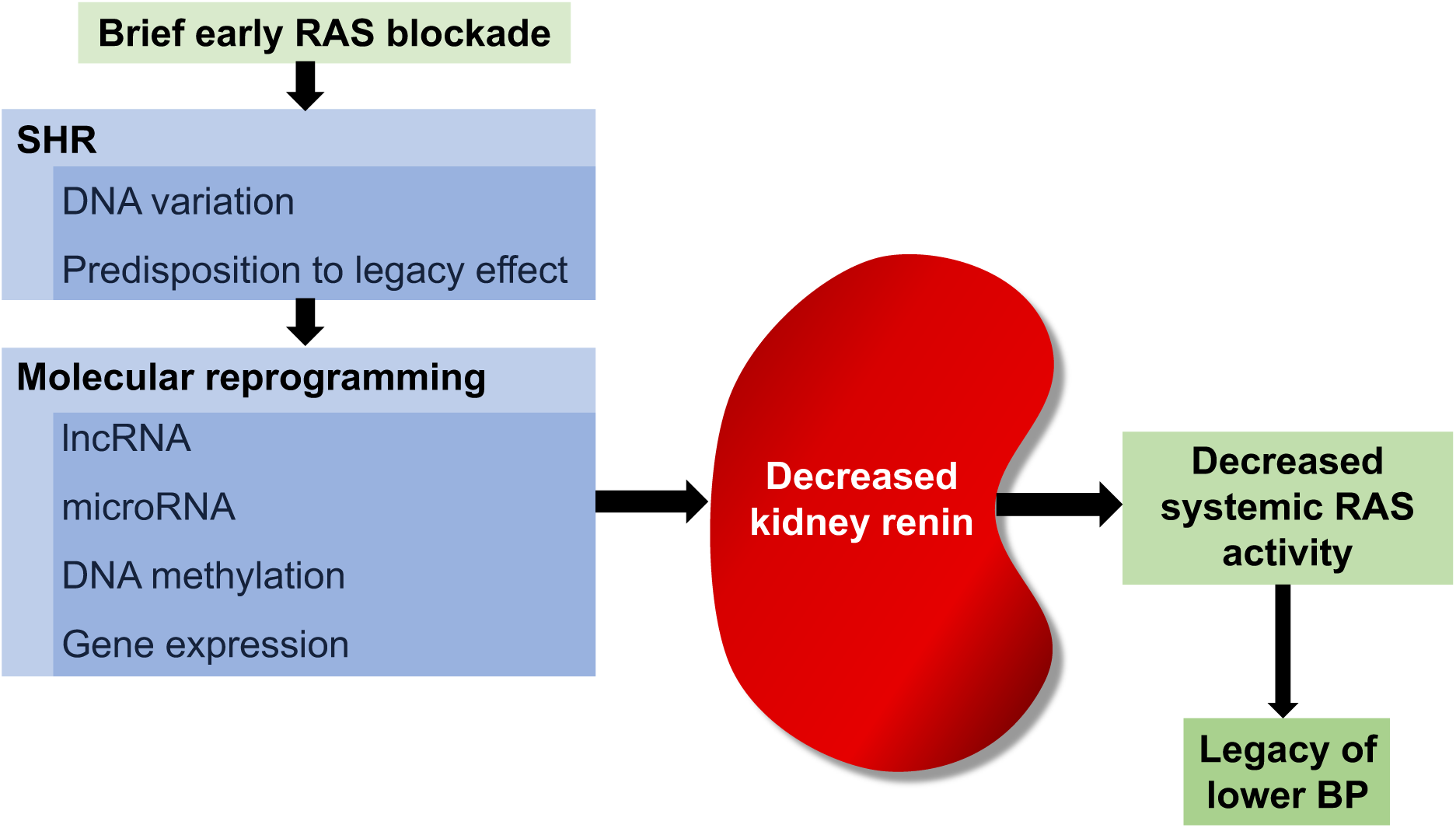

### Renin gene sequence variants

The details of 112 variants identified in and around the SHR-derived strains (not observed in BN or WKY) are shown in Table S18. These included SNPs, nucleotide repeat variants and insertion and deletions. Fifty-six of the variants were associated with changes in the sequence of putative transcription factor binding sites including a previously reported variant^38^ in the first intron for GR-alpha. Several of the transcription factors already have established relationships with the control of renin expression and the RAS including FOXP3,^39^ RXR-alpha,^40^ PPAR-alpha,^40^ RBP-J^41^ and c-Jun.^42^ Three sequence variants in the antisense lncRNA LOC102550525 represented changes to putative transcription factors binding sites (Table S18).

## DISCUSSION

Since the initial observation some 40 years ago,^2^ numerous independent studies in SHR have established that brief RAS inhibition in younger animals can prevent the full development of genetic hypertension. Despite considerable effort, the explanation for this remarkable phenomenon has remained elusive. Here we report for the first time that following early ARB or ACE inhibitor treatment in SHR the persistent reduction in BP, instead of eliciting the expected reflex increase in renin^43^, is associated with a reduction in kidney renin, as evidenced by significantly lower gene expression and protein immunolabelling.

That the lower kidney renin should be important to the legacy of early RAS inhibition in SHR is substantiated by the major role of the RAS in BP homeostasis generally and by the pathophysiology of SHR hypertension specifically. Renin drives the RAS and its control of BP through regulation of the initial and rate-limiting step of the RAS cascade. The kidney is the only organ capable of releasing enzymatically active renin.^44^ The kidneys and renin are also relevant to the pathogenesis of SHR hypertension. High plasma and kidney renin activity^19, 20^ and abnormalities of renal glomerular hemodynamics are seen during the development of SHR hypertension.^45^ Our previous crossbreeding studies have genetically linked BP, glomerular hemodynamics, and plasma renin activity to the development of hypertension.^21^ The involvement of kidney renin with the legacy effect also aligns with two other important observations. The first is that RAS inhibitors, but not other classes of antihypertensive drugs, produce a long-term reduction in BP. The second is that the kidneys are central to the legacy effect,^46^ as epitomised by transplantation in which the BP “follows the kidney”.

Surprisingly, the potential involvement of the RAS in long-term BP effects of RAS inhibition in SHR has received relatively little attention. Previous studies revealed long-term reductions in plasma angiotensinogen levels after early ACE inhibition^47^ and reduced renal angiotensin II and aldosterone,^4^ but kidney renin was not measured. Few data from other studies are available, but in the closely related SHRSP strain, the development of severe renal damage as a result of advance age^7^ or L-NAME treatment^12^ caused an increase in kidney *Ren* expression but this response was significantly attenuated in SHRSP that had received earlier treatment with the ARB candesartan, suggesting long-term resetting of kidney renin.

Why might kidney renin remain low after early RAS inhibition in SHR? One physiological explanation could be reduced sympathetically-mediated renin release.^44^ We have no direct measure of sympathetic activity, but the heart rate measured by telemetry in freely moving SHR treated with either losartan or perindopril was not different to controls either during or after treatment (data not shown). Furthermore, transplantation studies^23^ in which renal nerves are severed suggest that factors other than renal sympathetic activity explain the long-term lower BP.

Instead, we believe that the reduced kidney renin is the result of genetic reprogramming by early RAS inhibition, a phenomenon to which the SHR is genetically predisposed (Figure 7). Our genomic analyses provided insights into the nature of this reprogramming. We found significant changes in expression of genes, non-coding RNA and DNA methylation. Our network analyses revealed changes in expression or methylation that showed strong links with the reduced *Ren* expression. Furthermore, the gene ontology analyses indicated that the reprogramming involved genes with known relationships with RAS, BP, and the kidneys. These networks included 13 candidate genes that were differentially expressed at 20 weeks of age after early RAS inhibition and that we believe are likely to cause or sustain the reduced *Ren* expression and the legacy of lower BP.

Among the 13 candidates, six (*Grhl1*, *Nfil3*, *Adamts1*, *Hs6st1*, *Adap1*, *Bcl6*) have established cardiovascular involvement and interactions with the RAS and nine showed differential expression in the same direction after perindopril and losartan, possibly reflecting class effects of RAS inhibition. Within these candidates, *Nfil3* is of special interest. Only *Nfil3* showed significant evidence of reduced expression during (at 14 weeks in LOS and at 20 weeks in PER_late_) and after (20 weeks in LOS and PER) RAS inhibition treatment. There was also evidence for significantly increased methylation of *Nfil3* at 20 weeks of age (Table S16). In addition, *Nfil3* was a hub gene in a co-expression module (purple, Table S3) that also contained the greatest number of candidate genes (*Grhl1*, *Ammecr1l*, *Hs6st1*, *Nfil3*, *Fam221a*, *Lmo4*, *Adamts1*).

NFIL3 (nuclear factor, interleukin 3 regulated, also known as E4BP4) is a transcription factor involved in control of circadian rhythm that is associated with BP and has links to the RAS. Circadian clock genes have been identified as abnormally expressed in the kidneys of SHR^48^ and *Nfil3* is over-expressed in SHR adrenal glands.^49^ Conversely, *Nfil3* knockout mice have reduced systemic BP.^50^ In terms of interaction with the RAS, we observed significant reduction in *Nfil3* expression during RAS inhibition. Previous studies have demonstrated that NFIL3 has potent effects to increase expression of aldosterone synthase and aldosterone secretion, an effect that is amplified by angiotensin.^51^ Furthermore, *Nfil3* knockout results in the loss of angiopoietin-2,^50, 52^ while the expression of angiopoietin-2 is reduced by angiotensin receptor antagonism.^53^ Therefore, reduced *Nfil3* expression as a result of RAS inhibition could further amplify the suppression of the RAS. Interestingly, recent genome-wide meta-analysis also identified *Nfil3* as significant potential gene target for hypertension.^54^

Of the other candidates we identified, *Adamts1* (ADAM metallopeptidase with thrombospondin type 1 motif, 1) also showed significantly reduced renal expression at 20 weeks of age. ADAMTS1 is a metalloproteinase with a number of functions that include inhibition of angiogenesis,^55^ blood vessel remodelling^56^ and the normal development of the kidney.^57^ Importantly, angiotensin II induces expression of *Adamts1* in endothelial and vascular smooth muscle cells.^58^ Mice with a genetic haploinsufficiency of *Adamts1* have significantly lower BP than controls.^59^

In contrast, the expression of *Bcl6* was significantly increased in the 20-week LOS SHR. BCL6 is a sequence-specific transcriptional repressor that shows increased renal expression in young SHR^60^ but lower expression as hypertension develops in adulthood,^61^ as we also observed in vehicle SHR (Figure 3). Other studies have shown that over-expression of *Bcl6* significantly reduces SHR BP.^61^ At a mechanistic level, BCL6 can inhibit angiotensin II-induced proliferation of human vascular smooth muscle cells (VSMC) and can attenuate angiotensin II-induced oxidative damage in VSMC and human renal tubular epithelial cells.^61^ Of special note was our observation that renal *Bcl6* expression showed the strongest negative correlation with *Ren* expression at 20 weeks.

Because of their regulatory effects on gene expression^24^ we specifically investigated miRNA expression and identified 45 miRNA molecules that were significantly differentially expressed at 20 weeks following losartan treatment (Table 2). Among these the miRNA miR-145-3p^62^ was strongly negatively correlated (r < -0.8) with nine out of the 13 candidate mRNA genes. miRNA-145 has been shown to be associated with SHR hypertension^63, 64^ and our observations suggest that it might play a significant role in the legacy effects following RAS inhibition.

To help understand how the identified candidate genes might influence the RAS, our multivariate analyses showed that seven of the candidate genes were highly correlated (|r| ≥ 0.8) with *Ren* at 20 weeks of age (Table S13). In relation to miRNAs that might be involved in the suppression of Ren expression, miR-145-3p showed the highest negative correlation (r = -0.77, Table S15). We also found links between lncRNA and miRNA expression at 20 weeks of age. In particular, there was a strong correlation (r = 0.92) between miRNA-145-3p and the lncRNA AC115371.3 (Table S11). This lncRNA showed the greatest number of correlations with the 45 top differentially expressed miRNAs (Table S11) and it was also highly negatively correlated (r = -0.82) with *Ren* expression (Table S10).

These gene network changes following early RAS inhibition in SHR provide insights into the molecular architecture underlying long-term genetic reprogramming. The origins of this reprogramming are initiated earlier during the active treatment period, where the key seems to be RAS inhibition in addition to lowering BP. We demonstrated major perturbations in the expression of elements of the RAS during active RAS inhibitor treatment, including a six-fold increase in *Ren* expression with increased expression of angiotensinogen, angiotensin II receptor associated protein and transforming growth factor beta 1. Such RAS gene over-expression may evoke compensatory homeostatic responses at a molecular level that have the effect of resetting genomic networks that impact on the RAS and the kidney, leading to reduced *Ren* expression. It is worth noting that long-term reduction in SHR BP depends on a sufficient period of treatment and is most effective in younger animals,^2^ suggesting that duration and timing of treatment influences the reprogramming set-points for *Ren* and cognate gene expression.

An important question is why reprogramming should persist after treatment stops. Epigenetic factors such as DNA methylation, histone modification and chromatin remodelling are relevant here. DNA methylation of CpG islands that are commonly found in promoter regions of genes^65^ characteristically occurs in a tissue-specific manner and can result in long-term effects by recruiting proteins involved in gene repression or by inhibiting the binding of transcription factor(s) to DNA. In this regard, our finding of increased methylation in the promoter region of *Ren* suggests a likely explanation for long-term reduced *Ren* expression.

In understanding the legacy of RAS inhibition, we also sought to find clues to the SHR (and SHRSP) should be genetically predisposed to this effect. The likely explanation is differences in DNA sequence variation, and we focussed on *Ren*. Based on the most comprehensive and accurate sequences available^29, 30^ we identified 112 variants in and around *Ren* that were found in SHR and SHRSP but not WKY or BN strains. Further experiments will be required to determine the potential role these variants may play in predisposition to the legacy effects of RAS inhibition. Of highest priority would be variants that may affect gene expression. In this respect we identified numerous variants in putative binding motifs for transcription factors, many of which have been implicated previously in aspects of control of *Ren* expression.^38–41, 44, 66^ The impact of variants in the lncRNA LOC102550525 (antisense to *Ren*) also merit special attention given their diverse roles in regulation of gene expression.^67^

The overall picture emerging from these analyses is that the genomic landscape of renal expression during the legacy phase after RAS inhibition is a network of changes that involve the RAS itself. It would seem reasonable to attribute the long-term BP reduction to the lower renal *Ren* expression, particularly given that the kidneys are the primary source of renin, and renin is the rate-limiting factor in the RAS. It might also explain why the legacy BP effect follows the kidney in renal transplantation experiments. However, within the network of differential expression that we have been able to resolve, it is difficult to postulate with certainty the sequence of events or the hierarchy of control that results in the reduced *Ren* expression and persistently lower BP. However, our results provide the foundation for further experiments to test the specific contributions of individual candidates. Such experiments also have potential to define novel and targeted therapeutic approaches to replicating the legacy effect. Such information will be critical in informing future human trials in the prevention of hypertension.

### Novelty and Significance

What is known?

- SHR hypertension is genetically programmed and there is evidence of involvement of the kidneys and the RAS.
- Brief treatment with RAS inhibitors before adulthood results in a legacy of lifelong lower BP in SHR.
- The legacy BP effect “follows the kidneys” but the precise renal molecular mechanisms are unclear.

What new information does this article contribute?

- The legacy BP effect is associated with reduced renin gene expression and renin protein in the renal cortex.
- We detected associated relevant changes in candidate genes and miRNA and lncRNA expression and DNA methylation with influence on the RAS.
- DNA sequence variants in the SHR renin gene that might predispose to the legacy effect were identified.

### Summary

Despite the evident possibility, the involvement of kidney renin in the long-term legacy of reduced BP after early brief RAS inhibition in SHR has never been examined. Our comprehensive analyses for the first time reveal a reduction in kidney renin that provides a credible and logical explanation for persistently reduced BP. We have also defined the components of the molecular architecture that underpins the low renin. This is manifest as reprogramming of mRNAs, miRNAs and lncRNAs in genetic networks with known impact on BP, the kidney and the RAS, as well as evidence of epigenetic influences through increased methylation of the renin gene and other key genes. We also catalogued the DNA variants in and around the SHR renin gene that might predispose this strain to the long-term legacy effects after RAS inhibition. The added significance of these findings is foundational as they provide a basis for further experiments to investigate precise genetic hypotheses that have the potential to define new therapeutic targets to achieve the legacy effect with the possibility of targeted human trials among predisposed individuals.

## Acknowledgements

We thank Institut de Recherches Internationales Servier for the generous donation of the perindopril used in these experiments.

## Supplementary Materials and Methods

## Methods

### Telemetric direct arterial pressure recordings

Long-term continuous recordings of mean arterial pressure (MAP) and heart rate (HR) in conscious state were performed using a radiotelemetry system (DSI, St.Paul, MN, USA). Briefly, the rats were lightly anesthetised by inhalation of 2% isoflurane (ISOFLOTM, Zoetis, Australia) in an induction box prior to intramuscular injection of a mixture of ketamine (60 mg/kg, i.m.; Lyppard, Dingley, Australia) and medetomidine (250 µg/kg, i.m.; Pfizer Animal Health, West Ryde, Australia). Analgesia was provided by injection of a non-steroidal, anti-inflammatory agent (meloxicam, 1 mg/kg, s.c., Metacam, Boehringer Ingelheim, Sydney, Australia). The surgery started after loss of pedal withdrawal and corneal reflexes indicating the required deep surgical level of anesthesia. The eye’s moisture was maintained by gel application (Polyvisc). The surgical field was shaved and disinfected with 80% ethanol, chlorhexidine and betadine antiseptic solution. An incision in the midline of the abdominal cavity was made to enable isolation of the abdominal aorta caudal to the renal arteries. The catheter of the telemetry transmitter (TA11PAC40, DSI, St.Paul, MN, USA) was inserted into the abdominal aorta following the manufacturer’s instructions. At the end of the surgery, the animals were injected with warmed Hartmann’s solution (1 mL, i.p.; Baxter Healthcare, Australia), and the anesthesia was reversed with atipamazole (1 mg/kg, i.m.; Pfizer). Rats were placed on a heat mat for recovery, and then returned to their standard cage. The animals were monitored daily and allowed to completely recover for 10 days, before the recordings started. Both body weight assessment and 24 h BP recordings were performed once a week in each rat for 11 weeks. The BP recordings were sampled in 10 s segments with an acquisition frequency of 1000 Hz. The data summarized in the manuscript are an average from a 2h recording period during the dark (between 3:45-5:45 am) period.

### Tissue collection and RNA extraction

Animals were rapidly euthanized using isoflurane and ketamine and the kidneys were dissected on ice to obtain cortical tissue. All tissues were immediately submerged in RNA*later* stabilisation solution (Thermo Fisher Scientific), frozen in liquid nitrogen and stored at -80°C. Total RNA was extracted from dissected renal cortices from SHR vehicle (VEH), losartan (LOS) and hydralazine (HYD) treated animals using the miRVana™ miRNA isolation kit (Thermo Fisher Scientific) according to manufacturer’s instructions.

### Total RNA and miRNA sequencing, read processing and differential gene expression

Total RNA-sequencing, specific miRNA and methylation sequencing was performed by the Australian Genome Research Facility (Melbourne, Australia). Total RNA sequencing was also obtained from Novogene (Beijing, China) for certain groups including confirmation of VEH and LOS differential expression results. A total of 3 sequencing runs by AGRF and Novogene were made to accommodate all samples and differential expression analyses were made only by comparison within individual runs to avoid potential batch effects. RNA-sequencing was performed on samples from n=9 VEH (n=5 at 14 weeks and n=4 at 20 weeks of age), n=10 LOS (n=5 at 14 and n=5 at 20 weeks of age), n=3 HYD (20 weeks of age), n=5 PER (20 weeks of age).

All total RNA analyses utilized the same parameters including rRNA removal (Ribo-Zero depletion) and Illumina HiSeq 150bp paired-end sequencing at high depth (∼100 million reads per sample). Read quality was assessed using the FastQC software version 0.11.8 (www.bioinformatics.babraham.ac.uk/projects/fastqc/) and samples all showed highly quality base scores. Alignment and quantification of total RNA-sequencing data was performed using Rsubread aligner (version 1.34.6). Paired-end 150bp Illumina reads were aligned to the rat genome (UCSC rn6 assembly) at the gene level with an average of 97.6% (SD 0.85%) of reads successfully mapped. Genes with greater than one count-per-million mapped reads in at least two samples were retained for further analysis, genes below this threshold were filtered out.

miRNA single-end 50bp Illumina reads were quality checked using FastQC version 0.11.8 (www.bioinformatics.babraham.ac.uk/projects/fastqc/) and samples all showed highly quality base scores. miRNA reads were aligned and quantified with Oasis 2.0,^1^ an online software package specialised for small RNA-sequencing data including trimming of the Illumina adapter sequence from fastq files, read filtering (15–32 nt) and removal of low abundance reads (< 5 reads). Remaining reads were mapped to the rat genome rn6 and miRBase v22 with an average of 75.6% (±10.2 SD) of reads successfully mapped.

Analysis of differential expression in total RNA and miRNA sequencing data was performed in the R statistical programming environment (version 3.5.2) using edgeR (version 3.26.7), EDAseq (version 2.18.0) and RUVseq (version 1.18.0) Bioconductor packages. Biological coefficient of variation (BCV) was checked using the common dispersion method (negative binomial dispersion by conditional maximum likelihood). BCV is derived by subtracting the estimated technical variation (i.e. measurement error) from total CV across libraries (see edgeR manual available on Bioconductor.^2^

Data normalisation including adjusting for library size and removal of potential batch/technical effects was performed using EDAseq and RUVseq (Remove Unwanted Variation from RNA-Seq data). The function betweenLaneNormalisation in EDAseq was used for sequencing depth normalisation among samples using a nonlinear full quantile method.^3^ A second normalisation step based on factor analysis of putative non-differentially expressed genes was performed. This included producing a set of *in silico* empirical control genes (undifferentiated FDR q > 0.95) via differential expression analysis in edgeR.^4^ These negative control genes were used in the RUVg normalising function in RUVseq,^2^ removing k=2 factors of unwanted variation. Normalization success was checked by plotting the relative log expression (plotRLE function) and the PCA (plotPCA function) across samples using EDAseq.

Differential expression analysis was performed in edgeR, which applies a trimmed mean of M values (TMM) normalisation where dispersion parameters for each gene are estimated with the Cox-Reid common dispersion method^5^ and employed in a negative binomial generalized linear model for each gene. Accounting for gene dispersion ensures that expression differences that are consistent between replicates are more highly weighted than those that are not to ensure differential expression is not driven by outliers. P-values were adjusted for multiple testing using the Benjamini-Hochberg correction with an FDR q-value < 0.05.

### Methylation sequencing and read processing

Illumina NovaSeq (50bp single-end reads, ∼10-20 million reads per sample) was used for methylation sequencing using the reduced representation bisulfite sequencing (RRBS) technique. RRBS single-end 100bp Illumina reads were processed using the following steps also described in the protocol by Chen *et al*.^6^ and the edgeR package user’s manual.^7^ Reads were first trimmed with Trim Galore version 0.6.6,^8^ a wrapper for cutadapt version 3.5,^9^ with the *-rrbs* flag set to remove adapters and trim poor quality reads. Trimmed reads were then aligned with Bismark version 0.22.3,^10^ to the rn6 rat genome using Bowtie2.

Methylation calls were made using the bismark_methylation_extraction function and read into R with the readBismark2DGE function in edgeR. CpGs on unassembled chromosomes and those assigned to the Y chromosome and mitochondrial DNA were removed, and CpGs were annotated with the identity of the nearest gene with the nearest transcription start site (TSS) function. CpGs with low coverage were filtered out by summing the counts of methylated and unmethylated reads to get the total read coverage at each CpG site for each sample. CpGs with a total count (methylated and unmethylated) of less than 8 in every sample were removed as well as CpGs that were never methylated or always methylated as they provide no information about differential methylation^5^.

### Genome wide differential methylation analyses for CpGs and gene promoters

The genome-wide analyses detected over 9 million CpGs from which we filtered out CpG loci with very low reads across samples. Testing for differential methylation was performed using the ratio between methylated and unmethylated counts modelled by a negative binomial linear model in edgeR. To assess methylation in gene promoters, CpG counts were aggregated 2kb upstream to 1kb downstream of the transcription start site (TSS) for all annotated genes. The gene promoter approach fits well with the observation that CpG methylation in promoter regions is often associated with silencing of transcription and gene expression.^11^ To assess methylation in CpG clusters, CpG counts were aggregated in naturally occurring CpG-rich islands (see methods for aggregation in section below). We did not interpret methylation at individual CpGs (i.e. at the highest possible resolution of methylation) due to variation in methylation status (methylated/unmethylated) per CpG across samples and differences in differential methylation often observed between directly adjacent CpGs (within islands) which often resulted from low/variable read numbers per CpG even after applying standard read count filters in edgeR.

### Genome wide differential methylation analysis for CpG clusters

In order to investigate patterns of genome wide methylation outside of gene promoter regions, we assessed methylation in CpG clusters (islands) with custom R code. This code was designed to scan across the entire genome and aggregate counts (same as applied in the gene promoter analysis) into naturally occurring CpG clusters – every time a gap between two CpGs of ≥500bp was encountered, a new CpG cluster was assigned. This was initially tested on a smaller subset of genes and matched well with CpGs that visually clustered within those genes. This genome-wide cluster-level approach fits with the finding that over half of all CpG islands are found outside of TSS/promoter regions.^12^

### Multivariate analyses in MixOmics to explore potential gene co-regulation between coding and non-coding genes at 20 weeks related to Losartan treatment

Given the importance of gene regulatory functions of non-coding RNA, we investigated multivariate correlations between differentially expressed (FDR q<0.05) coding and non-coding RNA in all datasets that included: 272 protein coding genes, 15 lincRNAs and 19 other non-coding RNAs identified from the total RNA-seq datasets (see Table S1); 45 miRNAs identified in the miRNA-seq dataset (see Table S2).

We used a multivariate framework which allows expression of all differentially-expressed coding and non-coding genes across LOS and VEH samples to be compared (correlated) simultaneously, while also employing hierarchical clustering (complete linkage) to group coding and non-coding genes with highly similar gene expression profiles which can be used to infer potential gene co-regulation. We employed regularized Canonical Correlation Analysis (rcc function) in the R package mixOmics version 6.6.2.^13^ to perform the multivariate correlations. These multivariate correlation patterns were then visualised with clustered image maps (cim function, mixOmics) from which network diagrams were constructed with only the most highly correlated (|r| ≥ 0.8) genes to form hypotheses about potential co-regulatory relationships between coding and non-coding genes.

#### Coding vs non-coding differentially expressed genes

The clustered image map in Figure S2A shows all differentially expressed coding and non-coding genes and provides a broad overview of gene clustering within and between coding and non-coding genes. Figure S2B which was derived from Figure S2A shows only coding and non-coding genes that were highly correlated (|r| ≥ 0.8) only – this network diagram was constructed in Cytoscape version 3.7.1. Figure S2C shows a simplified network diagram with only the 13 candidate genes.

#### Co-regulation between RAS genes and 35 confirmed mRNAs and 45 miRNAs

To explore potential co-regulation between RAS genes and the 35 confirmed genes at 14 and 20 weeks, as well as between RAS genes and miRNAs at 20 weeks, we performed multivariate analyses (same methods described above) between each dataset. Figure S5 (A-C) shows the clustered image maps and derived highly correlated (|r| ≥ 0.8) pairs only, shown in the network diagrams.

### Co-expression analyses in WGCNA to define gene networks at 20 weeks related to Losartan treatment and BP differences

We used WGCNA version 1.68,^14^ which aims to build a gene networks which are based on sets of modules consisting of highly interconnected genes with similar expression profiles. These can be used to analyse the internal relationship between key genes in modules and clinical characteristics. The main analysis steps are as follows: a correlation matrix between pairs of expressed genes is generated; hierarchical clustering is performed before constructing the co-expression network, and a soft thresholding power (β) is used to conform the relationship between genes in the gene co-expression network to the scale-free network distribution; the adjacency matrix is then converted into a topological overlap matrix (TOM), and the corresponding degree of difference (1-TOM) calculated; a minimum number of genes per module is defined and modules divided according to the standard of a dynamic shear tree; finally, module statistics (e.g. module eigengenes, module membership) are calculated and modules can be directly correlated with clinical characteristics.^15^

Three WGCNA networks were generated, one for each class of studied genes (mRNAs, miRNAs and non-protein coding RNAs such as lncRNAs and snoRNAs). In order to retain a sufficient number of genes for network construction in WGCNA yet still focus on genes related to treatment, we included any differentially expressed gene with a nominal p-value<0.05 from the total RNA-seq data (n=2844 protein coding, Table S3; 186 non-coding RNAs, Table S4). For the miRNA data, we chose a more liberal p threshold (p<0.10) to retain a sufficient number of genes for network construction (n=210 miRNAs, Table S5).

For each network, genes within modules were assessed for their intramodular connectivity (IC, connectivity of genes with other genes within the same module), module membership (MM, correlation of individual gene expression with module eigengene (ME) of its respective module) and hub genes. In scale-free network topology, ‘hubs’ are the most highly connected genes (of which there are typically relatively few) that correspond with high values for IC and MM. To identify gene modules (coding and non-coding) that might be related to the legacy effect on BP, we examined the correlation between ME and SBP (Tables S3-S5).

### Gene ontology enrichment analysis to examine biological pathways related to Losartan treatments effects and the RAS

To obtain an insights into potential functional relevance of differentially-expressed genes we undertook two gene ontology (GO) enrichment analyses with the Panther Classification System release 17.0 in the Gene Ontology (release 2023-05-10) application.^16, 17^ Analysis 1 included mRNA genes in modules (Table S3) that had been defined from WGCNA analyses, however there were often insufficient numbers of genes within modules to perform the overrepresentation tests in the GO analysis. Analysis 2 included leading differentially expressed genes related to LOS treatment (n=1676, Table S7). These analyses accounted for multiple testing (FDR) by applying the Benjamini-Hochberg procedure to Fisher’s exact test p-values.

### Renin immunolabelling

Immunohistochemistry for renin was performed as described previously^18, 19^ and all analyses were blinded to the treatment groups. Briefly, rat kidneys were fixed in 10% buffered formalin. Five μm paraffin sections of kidney were incubated with normal goat serum for 1 hour (5425S, Cell Signaling Technology), and then overnight at 4°C with a polyclonal mouse renin protein antibody raised against pure mouse submandibular gland renin (1:8000) (1. 2). A negative control without the primary antibody was included. Sections were washed with 0.1M phosphate buffered saline (PBS), incubated for 1 hour with biotin-conjugated goat anti-rabbit IgG (1:500, 111-065-144, Jackson ImmunoResearch, Pennsylvania, USA), washed with PBS and then incubated with the Vectastain ABC standard kit (Vector Laboratories, Pennsylvania, USA) for 30 minutes and liquid DAB substrate chromogen kit (BD Pharmingen, BD Biosciences, CA, USA) for 5 minutes. Following rinsing with tap water, sections were counterstained in Mayer’s hematoxylin and coverslipped. For quantitation, four randomly chosen sections from each kidney at least 125 μm apart were selected. Images of renin immunolabelling associated with the juxtaglomerular apparatus (JGA) from 50 glomeruli per section were captured at 100X magnification using a digital microscope camera (DS-Ri2, Nikon, Japan) attached to an upright microscope (H550L, Nikon, Japan. The fraction of JGAs that were renin positive was averaged across four sections from each sample.

## SUPPLEMENTARY FIGURES

**Figure S1:**
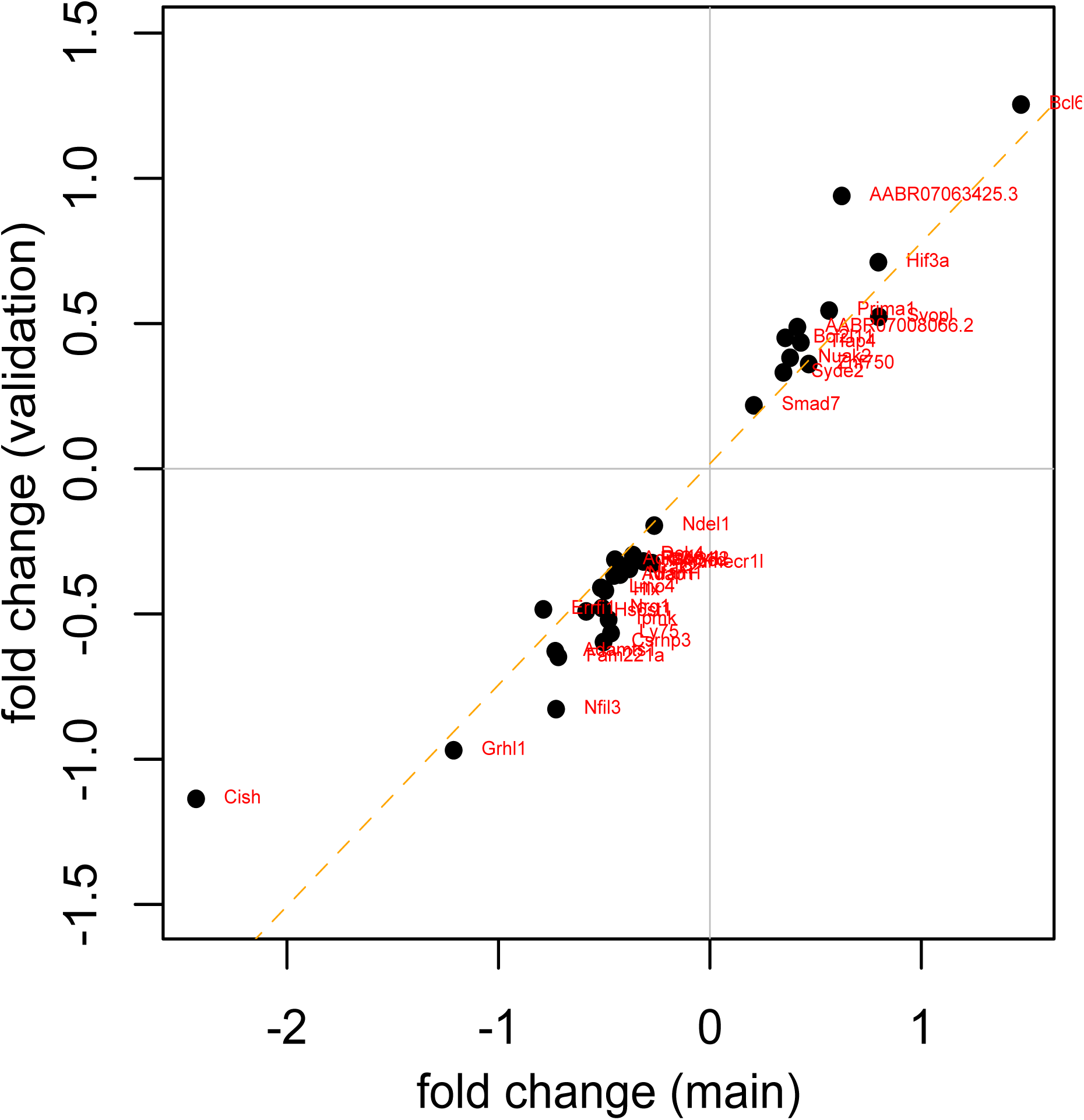
Comparison of log base 2 fold change (log_2_FC) expression for 35 genes between LOS and VEH SHR animals at 20 weeks. X-axis represents the initial (main AGRF) dataset, y-axis represents the validation (Novogene) dataset.

**Figure S2:**
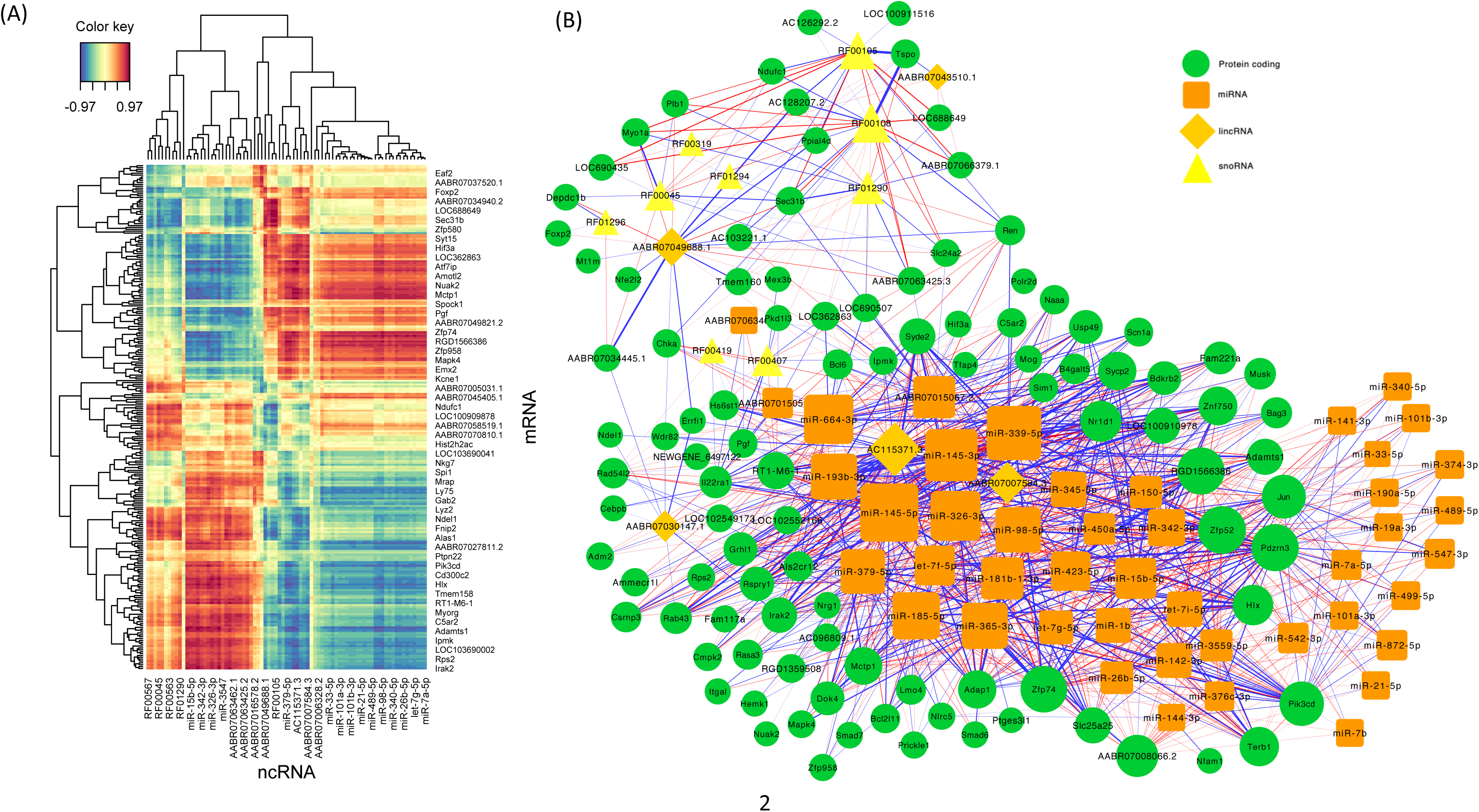
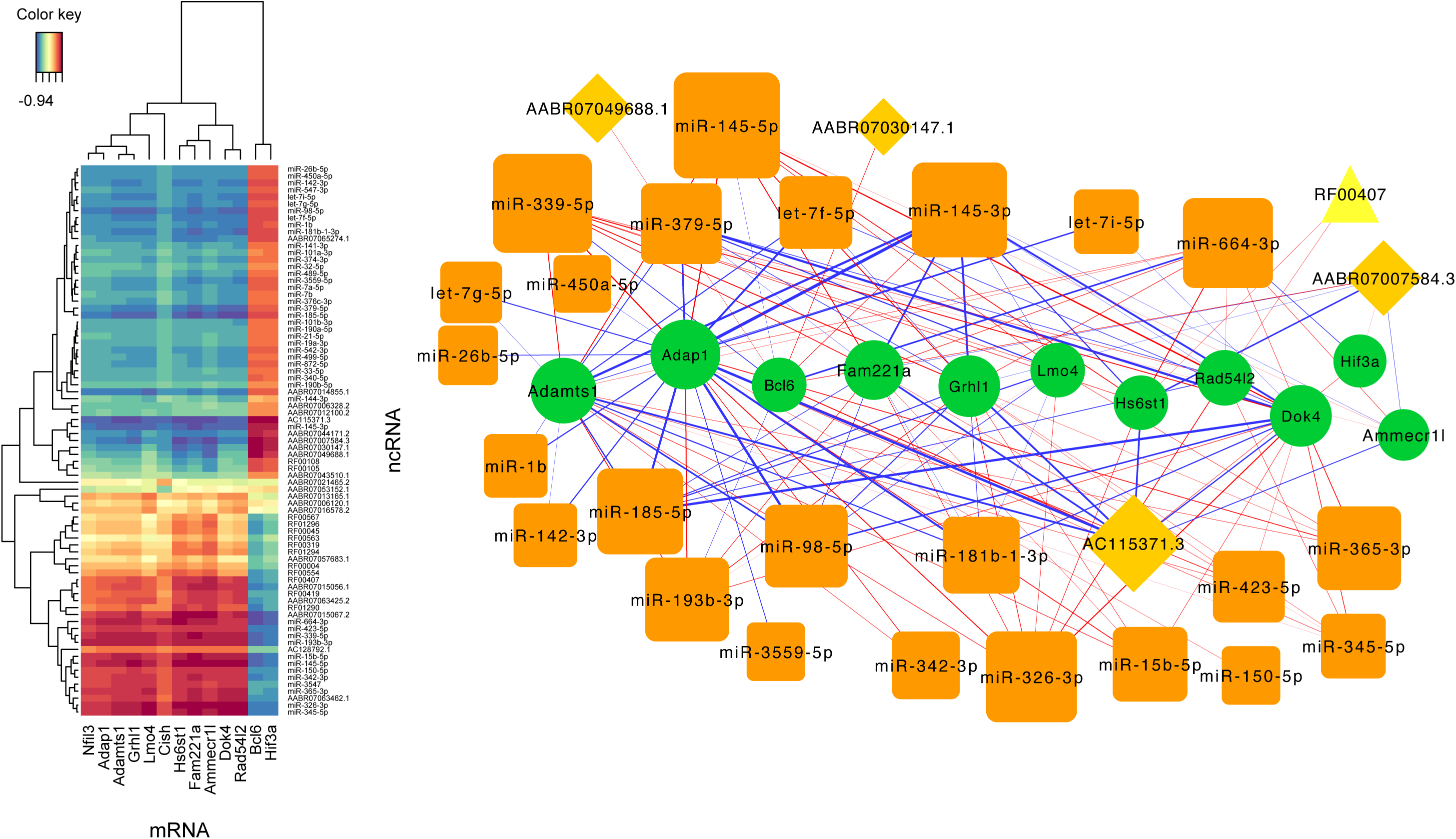
Multivariate correlations between differentially expressed coding and non-coding genes. (A) Clustered Image Map from canonical correlation analysis in MixOmics showing all correlations (see Table S10) and clustering patterns between coding (y-axis, not all gene labels shown) and non-coding (x-axis, not all gene labels shown) genes based on hierarchical clustering. (B) Cytoscape network between most highly correlated (|r| ≥ 0.8) coding and non-coding genes. Symbol size is determined by number of correlations (|r| ≥ 0.8) between coding and non-coding genes. Line weight determined by magnitude of correlation with negative correlations emphasized (Cytoscape edge width: r = -0.8 to -1.0 with widths 0.1 to 3 respectively; r = 0.8 to 1.0 with widths 0.05 to 1 respectively). (C) Simplified network map (derived from Fig S2A-B) with only the 13 candidate genes (Table 1A) shown. Cytoscape network (right) shows only most highly correlated (|r| ≥ 0.8) genes. Two candidate (*Nfil3, Cish*) genes are not shown as they were not highly correlated (|r| < 0.8) with any on-coding genes. Network line weight determined by magnitude of correlation with negative correlations emphasized (Cytoscape edge width: r = -0.8 to -1.0 with widths 0.1 to 3 respectively; r = 0.8 to 1.0 with widths 0.05 to 1 respectively).

**Figure S3:**
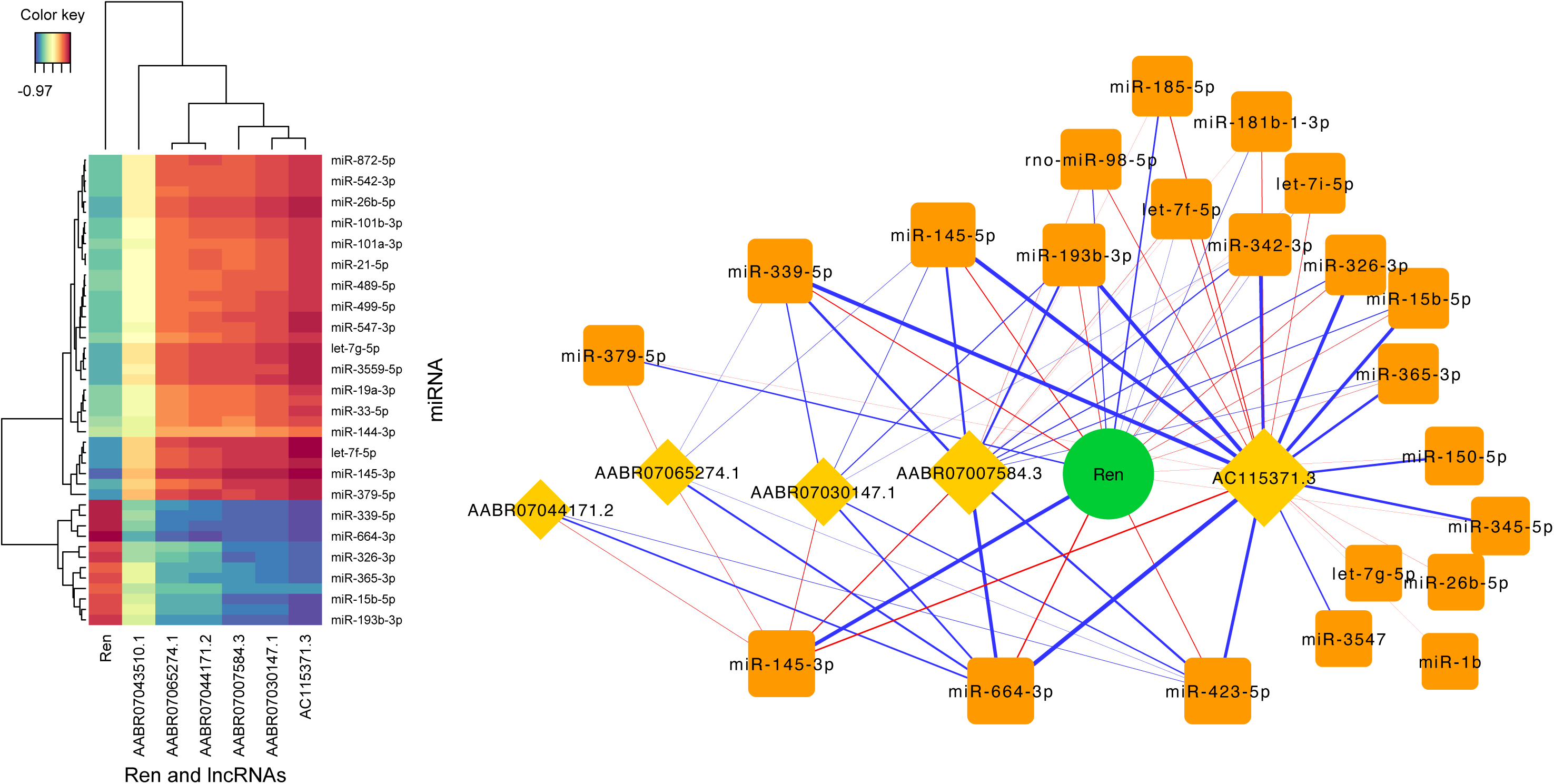
Multivariate correlations between differentially expressed lncRNAs, miRNAs and Ren. Clustered Image Map (left) from canonical correlation analysis in MixOmics showing all correlations and clustering patterns between miRNAs (y-axis, not all gene labels shown), Ren and lncRNA (x-axis) genes based on hierarchical clustering. Cytoscape network (right) between most highly correlated (|r| ≥ 0.8) genes. Symbol size is determined by number of correlations (|r| ≥ 0.8) between genes. Line weight determined by magnitude of correlation with negative correlations emphasized (Cytoscape edge width: r = -0.8 to -1.0 with widths 0.1 to 3 respectively; r = 0.8 to 1.0 with widths 0.05 to 1 respectively).

**Figure S4.**
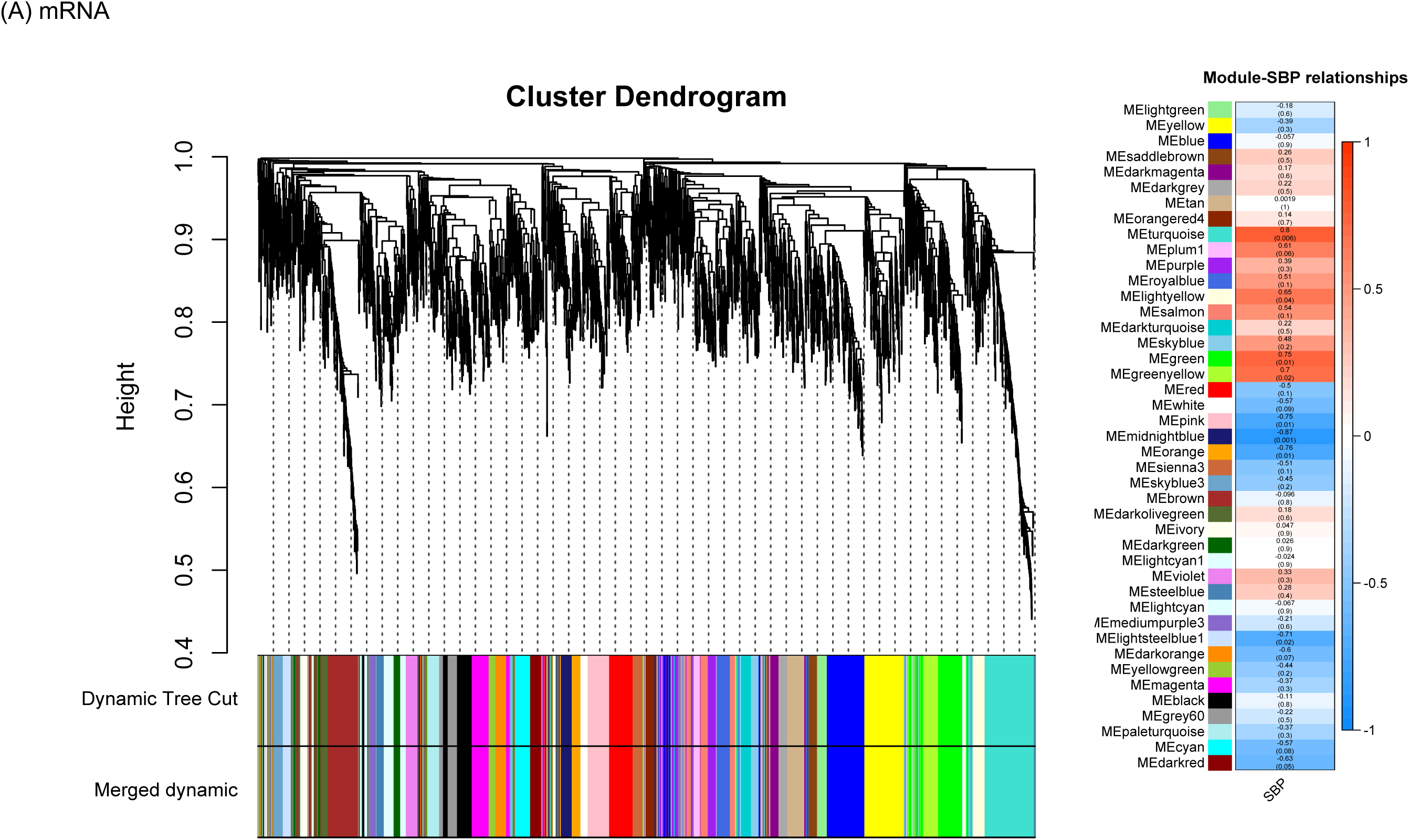
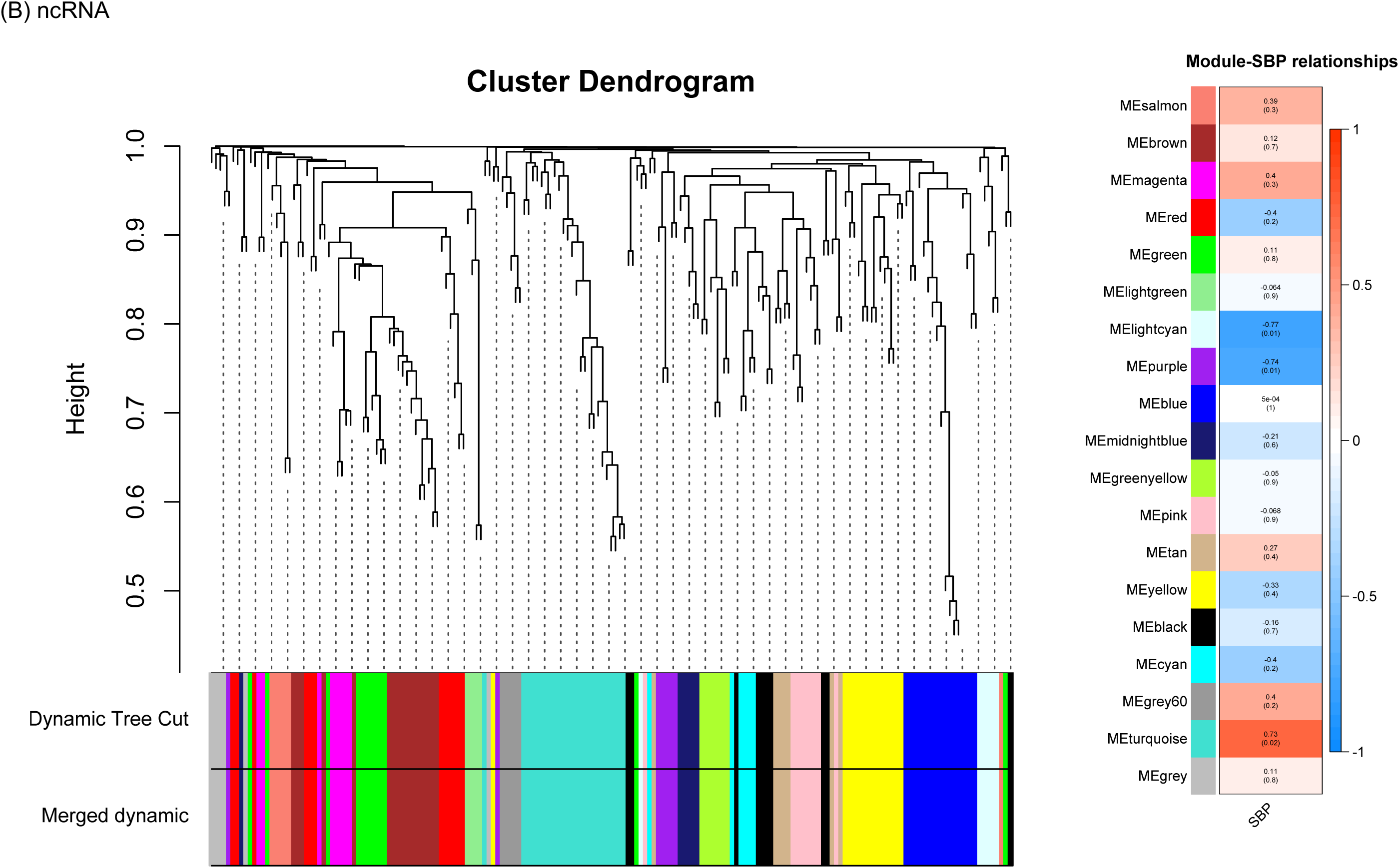
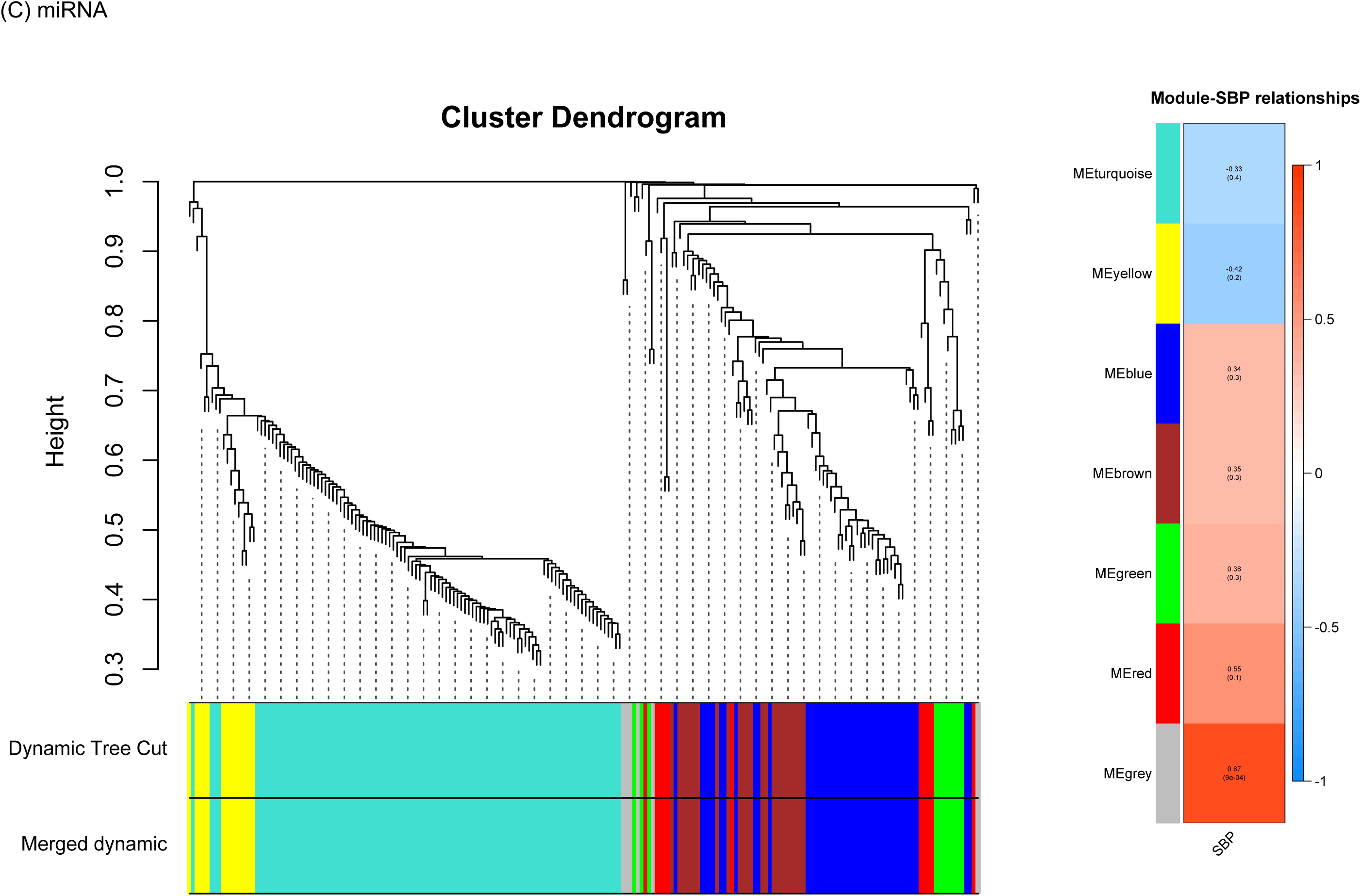
WGCNA analysis for mRNA (A), ncRNA (B) and miRNA (C). Left panel shows cluster dendrogram and module assignment, right panel includes correlation (upper values, vertical scale bar) and associated p-values (lower values) between module eigengene and systolic blood pressure measured at 20 weeks

**Figure S5.**
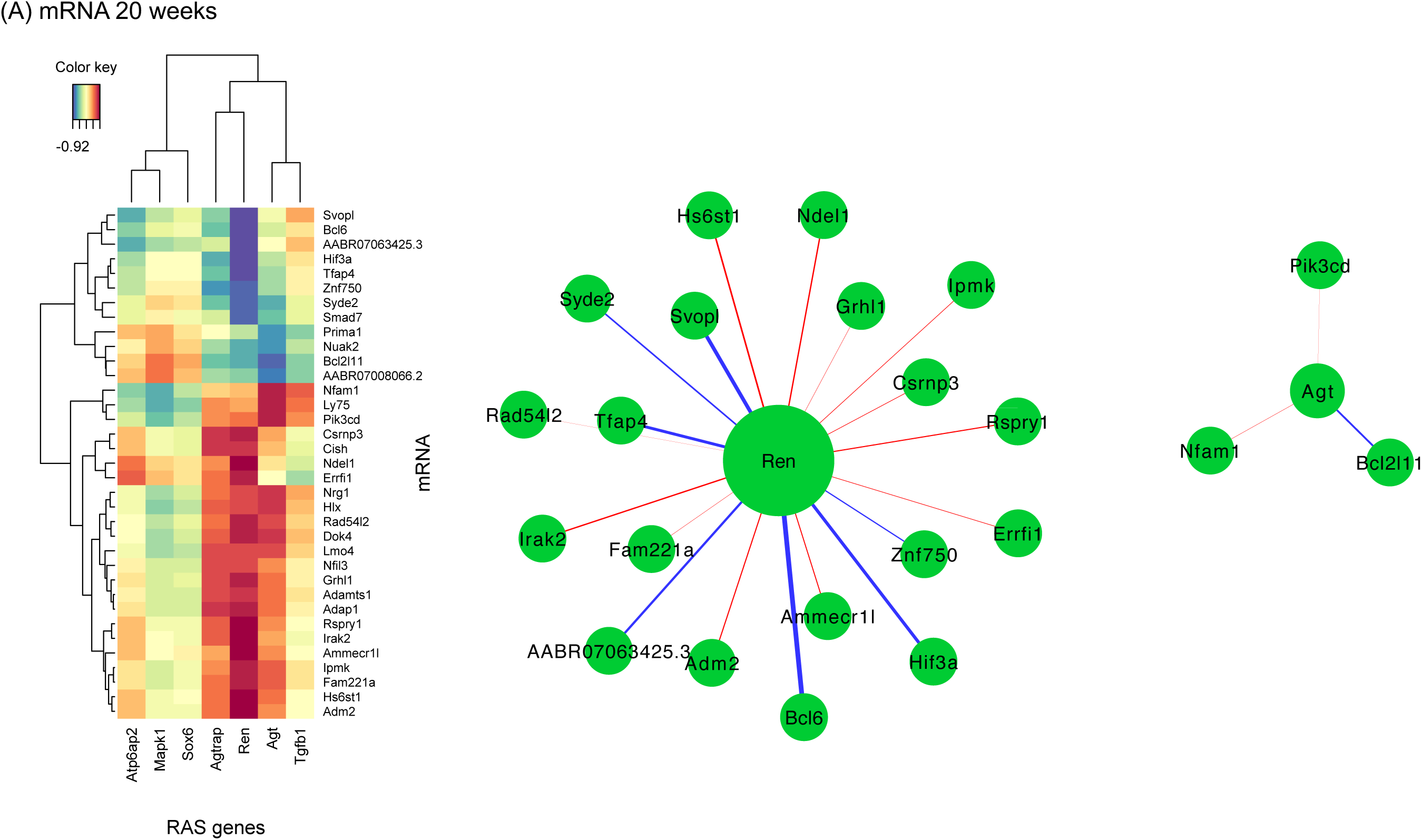
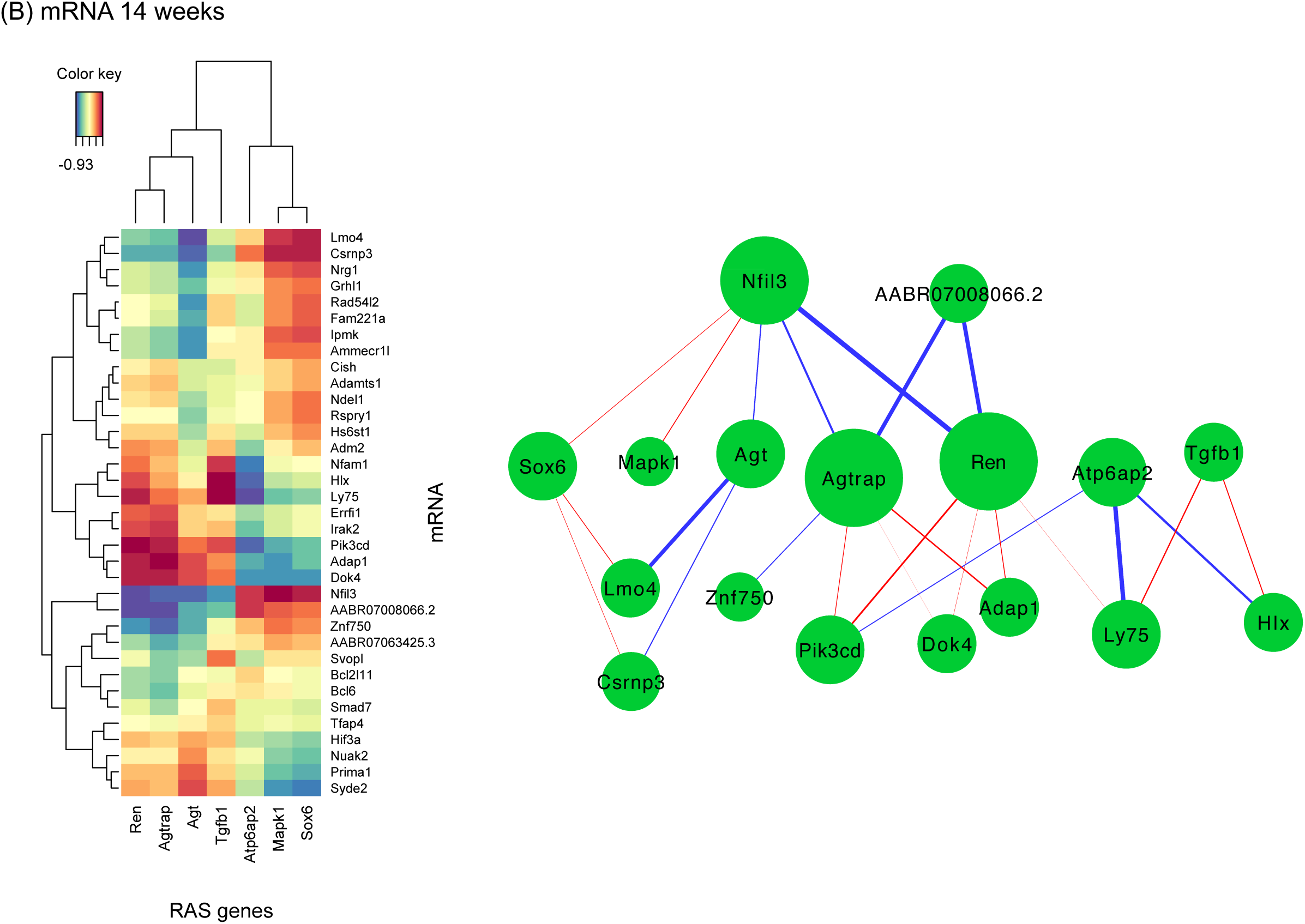
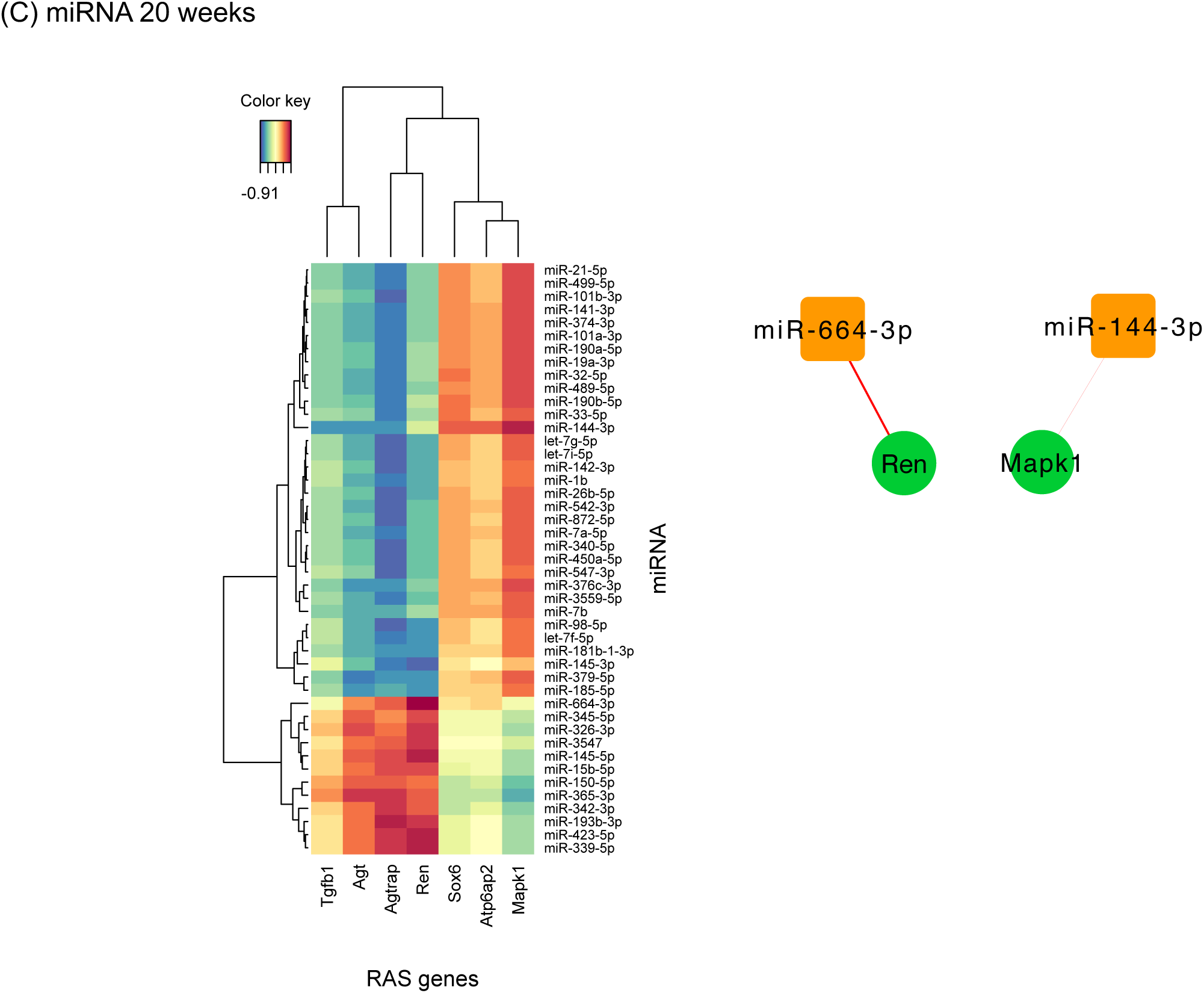
Clustered image maps and network diagrams based on correlations for RAS genes (|r| ≥ 0.8). (A) correlations at 20 weeks with validated 35 mRNA for differentially expressed genes identified at 20 weeks of age, (B) correlations at 14 weeks with validated 35 mRNA for differentially expressed genes identified at 20 weeks of age and (C) correlations at 20 weeks with top 45 differentially expressed miRNA identified at 20 weeks of age. Network line weight determined by magnitude of correlation with negative correlations emphasized (Cytoscape edge width: r = -0.8 to -1.0 with widths 0.1 to 3 respectively; r = 0.8 to 1.0 with widths 0.05 to 1 respectively).

